# Auto-inhibition of myoblast fusion by cyclic receptor signalling

**DOI:** 10.1101/553420

**Authors:** Daniel Sieiro, Julie Melendez, Valérie Morin, David Salgado, Christophe Marcelle

## Abstract

Fusion of nascent myoblasts to pre-existing myofibres is critical for skeletal muscle growth and repair. The vast majority of molecules known to regulate myoblast fusion are necessary in this process. Here we uncover, through high-throughput *in vitro* assays and *in vivo* studies in the chicken embryo, that TGFβ (SMAD2/3-dependent) signalling acts as a molecular brake on muscle fusion. While constitutive activation of the pathway arrests fusion, its inhibition leads to a striking over-fusion phenotype. This dynamic control of TGFβ signalling in the embryonic muscle relies on a unique receptor complementation mechanism, prompted by the merging of myoblasts with myofibres, each carrying one component of the heterodimer receptor complex. The competence of myofibres to fuse is restored through endocytic degradation of activated receptors. Altogether, this study shows that muscle fusion is a self-regulated process that relies on cyclic TGFβ signalling to regulate its pace.

## Main

Myoblast fusion occurs after muscle specification and early differentiation, themselves regulated by the Myogenic Regulatory Factors (MRFs) MYF5, MYOD and MYOG^1–3^. While MRF function is necessary for myoblast fusion^4^, terminal differentiation, including muscle contraction, can occur even if fusion is disrupted^5–7^. Work in *Drosophila* identified a number of molecules, mainly implicated in actin regulation, required for muscle fusion in this organism. A major advance was the demonstration that their homologs play similar function during vertebrate myogenesis^2,3,8^. Together with vertebrate-specific muscle fusion genes (e.g. Myomaker, Myomixer, JAM2-3) they constitute the “fusion machinery” necessary for the fusion of myoblasts to myofibres. Recent analyses we performed on myoblast fusion during embryonic development suggested that additional mechanisms must exert temporal and spatial control on the fusion machinery, modulating whether myoblasts and myofibres that are competent to fuse do so and at what pace^9^. The molecular underpinning of such control over fusion is completely unknown.

### A high throughput assay identifies novel regulator of myoblast fusion

To uncover those mechanisms, we performed an *in vitro* “esiRNA” (endonuclease-cleaved siRNA) screen on the mouse myogenic cell line C2C12. The esiRNA library represented about 9000 independent genes (i.e. one third of the mouse genome). This identified a large array of genes that either activate or inhibit C2C12 fusion, with little or no effect on differentiation and proliferation (Fig. 1a-c, Supplementary Fig. 1 and Supplementary Tables 1, 2). Validating the approach, we observed that molecules previously known to be necessary for myoblast fusion in vertebrates and invertebrates were identified through the screen (e.g. N-WASP, ARHGAP, MCADH, NFATC, IL4R, CAPN^10^, Supplementary Tables 1, 2 and not shown). These data constitute the first available resource for genes tested for fusion, proliferation and myogenic differentiation.

**Figure 1.**
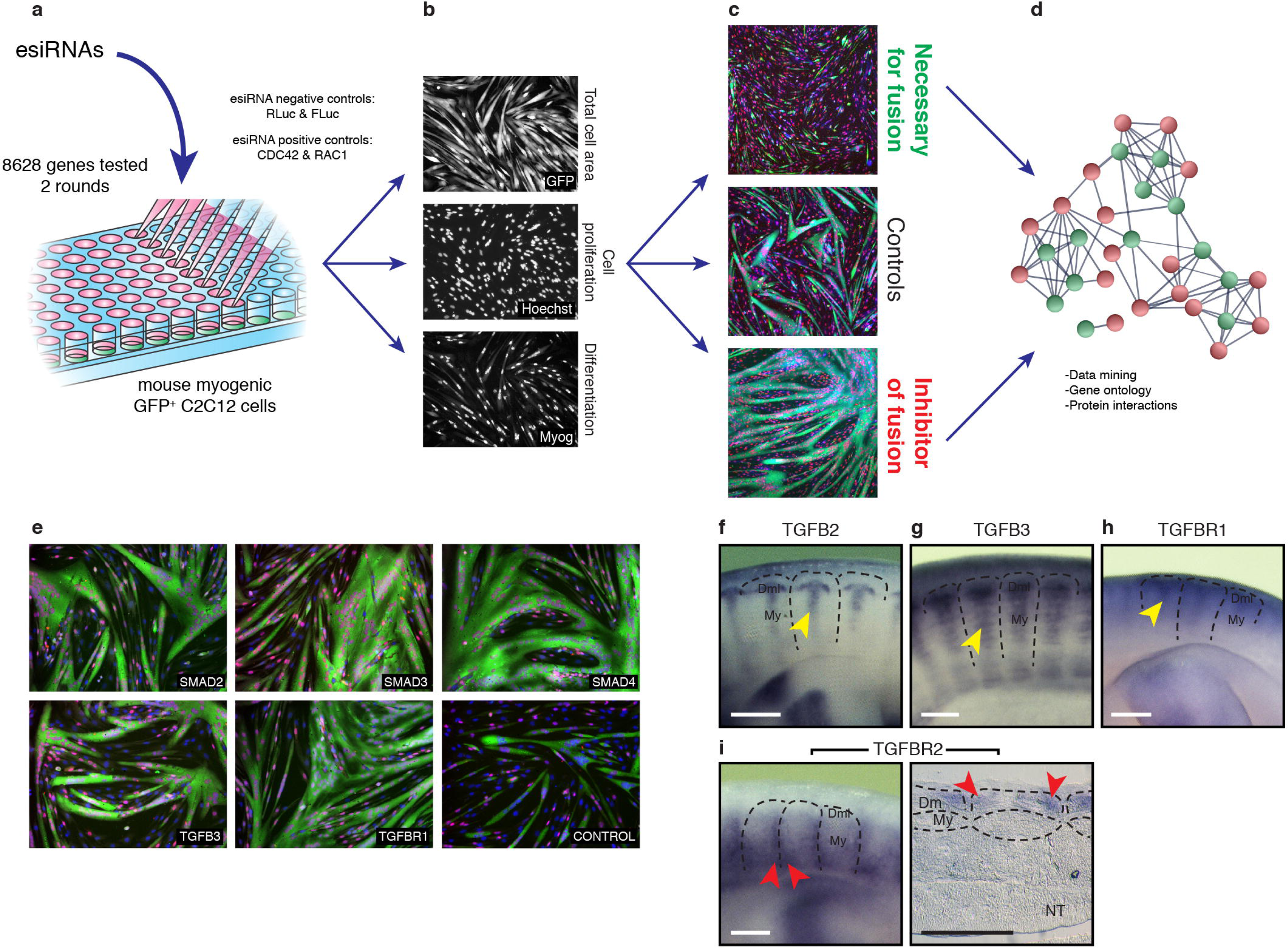
The TGFβ pathway acts as inhibitor of myoblast fusion *in vitro*, and members of the family are expressed in the muscle masses of the chicken embryo. **a**, EsiRNAs corresponding to 8628 distinct mouse genes were transfected into a (GFP-positive) C2C12 mouse muscle cell line to identify novel molecules regulating myoblast fusion. **b**, After differentiation, the area of myofibres (GFP staining), the nuclei count (Hoechst) and myogenin expression were evaluated with an image analysis program. **c**, Quantification of the number of nuclei per myofibre identified a number of genes that are either necessary or inhibit myoblast fusion *in vitro*, compared to controls. **d**, Gene ontology (Panther, Ingenuity) and protein interaction (String) analyses showed that the TGFβ pathway is over-represented within the inhibitors of fusion. **e**, Loss of function of ligands, receptors or effectors of the TGFβ pathway led to a strong increase of C2C12 fusion, compared to controls. **f-h**, Lateral view of whole mount (WM) *insitu* hybridization of chicken embryos at E5.5. The secreted ligands TGFB2 and TGFB3, as well as TGFBR1 are expressed in the myotome of the chicken embryo (yellow arrowheads, f-h). **i.** In contrast, TGFBR2 is expressed in epithelial cells located at the anterior and posterior borders of the dermomyotome (shown in WM and longitudinal section of chicken embryos at E5.5, red arrowheads). Dm: dermomyotome, My: myotome, NT: neural tube. Scale bars f-i: 0.5mm.

Gene ontology analysis software indicated that the TGFβ signalling pathway was over-represented within the group of molecules inhibiting the fusion of C2C12 cells (Fig. 1d, Supplementary Fig. 1 and Supplementary Table 3). Since inhibitors of fusion had not yet been identified, we sought to examine their role in this process *in vivo*. The TGFβ family is a large group of ligands that signal *via* heterotetrameric complexes of two type I and two type II receptors, which mediate their activity intra-cellularly via R-SMADs (Supplementary Fig. 2). TGFBs, activins, myostatin and nodal induce the phosphorylation of R-SMAD2 or 3, while BMPs mediate their activity through R-SMAD1, 5 or 8. All R-SMADs bind to SMAD4, an essential component for the generation of SMAD transcriptional responses^11–13^. Previous studies have demonstrated a role for some TGFβ ligands in myogenesis. While myostatin (MSTN) is best known for the dramatic increase of muscle mass that results from its loss-of-function, first identified in cattle^14^, it was also shown to promote the premature differentiation of resident muscle progenitors during embryogenesis^15^. In contrast, BMP represses early myogenic specification during development^16^. While these data suggest that BMPs and TGFBs play a role in the early steps of myogenesis (i.e. specification and differentiation), their role in the downstream events of the myogenic program (i.e. myoblast fusion) is still unknown.

In the screen, in addition to *bona fide* components of the TGFβ signalling pathway (TGFβ ligands -TGFBs-, receptors -TGFBRs- and SMAD effectors), we also identified positive (DPT, MMP14, RUNX1, SCUBE3) and negative (TGIF1) regulators of the pathway that promoted (TGIF1) or inhibited (all others) the fusion of C2C12 cells (Fig. 1e and Supplementary Table 3). Significantly, SMAD4, SMAD3, TGFB2 and TGFBR1 were among the 10% strongest inhibitors of fusion we identified in this screen. It is notable that none of the tested genes significantly influenced proliferation. While they displayed a mild inhibitory effect on myogenic differentiation (i.e. on Myogenin expression) as expected, the effect on fusion was in the majority of cases 2-3 times stronger than on differentiation, suggesting that their inhibitory activity on fusion was not solely due to their activity on upstream events of the myogenic program. Importantly, the members of the BMP signalling pathway and the Inhibins that were tested (BMP1, BMP4, BMP7, BMP8, GDF5, BMPR1A, ACVR2A, ACVR2B, SMAD5, INHBA, INHBC, INHBE) did not influence C2C12 fusion (Supplementary Table 3). These data suggest that TGFβ (but not BMP) signalling is a potent inhibitor of myoblast fusion *in vitro*.

### SMAD2/3-dependent TGFβ signalling acts as a molecular brake of myoblast fusion

We determined whether the TGFβ family members we identified were expressed at the time and place where we know that muscle fusion is taking place during embryonic development^9^. In the chicken embryo, trunk and limb muscle fibres initiate fusion at E4.5 (Stage HH25^17^). While the R-SMADs SMAD2 and SMAD3 are widely expressed in all embryonic tissues^18^, we found that the ligands TGFB2 and TGFB3 were specifically expressed in the myotome of somites from E4.5 (Fig. 1f, g) and in limb muscle masses one day later (not shown). At this stage of development, the myotome is composed of mononucleated, terminally differentiated myocytes (primitive myofibres) emanating from the medial border of the dermomyotome (the dorsal-most epithelial compartment of somites), named the DML^19–21^ (Supplementary Fig. 3). TGFB2 and 3 signal through TGFBR1 and TGFBR2^11^. TGFBR1 was expressed in the myotome of somites at E4.5 (Fig. 1h). Myocytes therefore express TGFB2 and 3 and TGFBR1. Remarkably, we observed that TGFBR2 was expressed at the anterior and posterior epithelial borders of each somite, but not in DML or in the myotome (Fig. 1i). This is important, since we recently demonstrated that myocytes derived from the DML exclusively fuse to progenitors originating from the anterior and posterior borders of somites^9^ (Supplementary Fig. 3). We performed gain and loss of function of TGFβ family members in developing somites. To avoid interfering with the proliferation, specification and early differentiation stages of myogenesis we used a myosin light chain (MLC) promoter^22^ to drive the expression of the various constructs we utilized. MLC is a gene activated late in myogenesis and coherent with this we observed that the activity of the promoter construct we used was initiated in terminally differentiating myogenic progenitors (Supplementary Fig. 4). Using the *in vivo* electroporation technique, we first over-expressed SMAD molecules under the control of the MLC promoter, co-electroporated with constructs coding for a membranal EGFP and a nuclear mCherry^9^, to evaluate the number of nuclei per electroporated myofibre. In this and the following experiments, when significant differences were observed we describe the changes in the proportion of fibres containing one nucleus, indicative of a tendency to arrest fusion, or more than 4 nuclei, indicative of a tendency to promote fusion. Solid colours in column graphs indicate significant differences while striped patterns indicate non-significant results.

Overexpression of SMAD4 by electroporation in the DML resulted in a significant (*** P-value<0.001) decrease in fusion in the population of DML-derived myocytes 3 days after electroporation (E5.5), at a time-point where fusion is largely under way throughout the myotome (Fig. 2a, b, f). In fact, a majority of electroporated myocytes remained mononucleated (+84%, compared to controls), while we observed a large deficit of fibres containing more than 4 nuclei (−57%, compared to controls). Similarly, overexpression of SMAD3 resulted in a significant decrease (*** P-value<0.001) in fusion when compared to controls (Fig. 2c,f). As with SMAD4, many SMAD3-expressing fibres remained mononucleated (+134%, compared to controls), while only few multinucleated fibres were observed (−96%, compared to controls). SMAD7 is a pan-inhibitor of TGFβ signalling (R-SMAD2/3-as well as R-SMAD1/5/8-dependent)^23,24^. Electroporation of SMAD7 in DML-derived myocytes led to a remarkable (*** P-value<0.001) over-fusion phenotype (Fig. 2d,f). Nearly all fibres had 4 or more nuclei (+512%, compared to controls) and only few were mononucleated (−91%, compared to controls). Moreover, we observed myocytes that contained up to 20 nuclei, while wild-type control myocytes contained 7 nuclei at most (Fig. 2 and not shown). Staining for Myosin Heavy Chain (MyHC) after over-expression of SMAD4, SMAD3 and SMAD7 (under MLC control) showed that all electroporated myocytes, regardless of their nuclei count, were terminally differentiated (Supplementary Fig. 5), suggesting that their expression did not interfere with the myogenic differentiation program. Even though BMP (R-SMAD1/5/8-dependent) signalling did not modify C2C12 fusion, we ruled out their implication in myoblast fusion *in vivo* by electroporating constitutively active SMAD1 and SMAD5 in DML cells. Three days later, the nuclei count of electroporated myofibres was not significantly different from controls (CA SMAD1, P-value: 0.16; CA SMAD5, P-value: 0.97; Fig. 2h-k). This confirms that BMP signalling does not play a significant role in the fusion of myoblasts to myofibres during early myogenesis. While the gain-of-function experiments described above indicate that TGFβ (R-SMAD2/3-dependent) signalling inhibits fusion *in vivo*, it was important to confirm these data by loss-of-function approaches, using the CRISPR-mediated gene-editing technique we recently applied to chicken embryos^25,26^. We targeted SMAD3 by electroporating the DML with a CRISPR/Cas9 vector containing two gRNAs directed against the chicken SMAD3 sequences. Results showed that loss-of-function of SMAD3 within muscle fibres significantly increased fusion (** P-value 0.002), compared to a CRISPR Cas9 vector containing no gRNAs (Fig. 2e, g), resulting in a decrease (−28%, compared to controls) in mononucleated fibres and an increase (+40%, compared to controls) in multinucleated myocytes. Altogether, these results demonstrate that, in somites, TGFβ signalling acts as a molecular brake on the fusion machinery. Inhibition of the pathway unleashes its restraint, resulting in a dramatic boost in the rate of DML-derived myofibres fusion, while its constitutive over-expression increases its constraint, leading to a near complete arrest of fusion.

**Figure 2.**
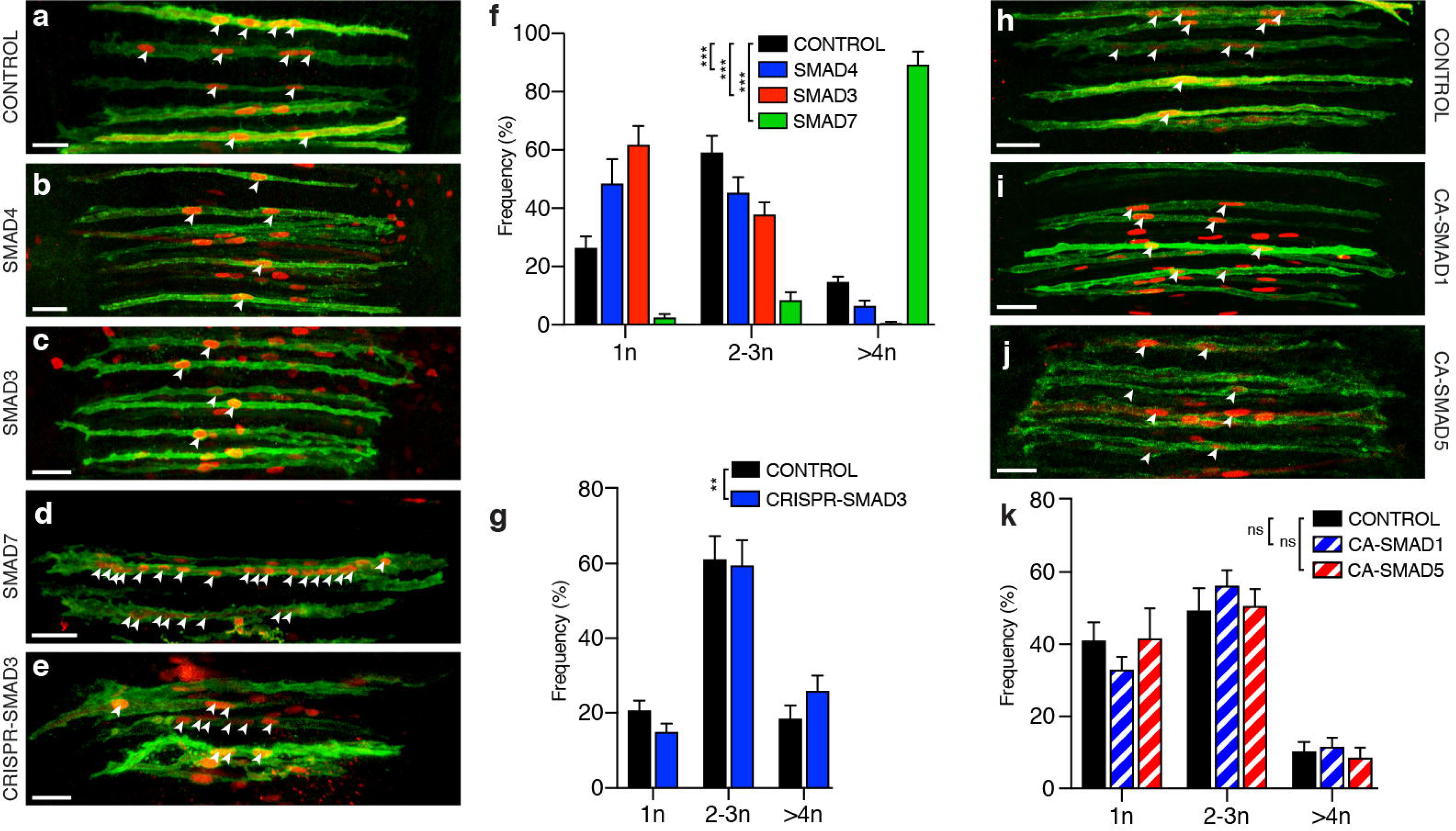
SMAD2/3-dependent TGFβ signalling regulates myoblast fusion *in vivo*. **a-d**, Dorsal view of confocal stacks of somites observed at E5.5. They were electroporated in the DML at E2.5 with MLC promoter driving expression of the indicated genes, together with membranal GFP (green) and nuclear mCherry (red) allowing the quantification of nuclei per myocyte. **e**, DML-derived myocyte observed at E5.5 and electroporated at E2.5 with a CRISPR/Cas9 construct targeting the chicken SMAD3 sequence together with membranal GFP (green) and nuclear mCherry (red). **f**, Column graph for a-d showing the population of electroporated myocytes distributed in those that contained one, two to three, and four or more nuclei, relative to its control (in %). Experimental treatments that led to significant differences compared to controls are indicated on all graphs by solid colour columns. **g**, Column graph for e. h-j DML-derived myocyte observed at E5.5 and electroporated at E2.5 with MLC promoter driving gene expression of the indicated BMP-related SMADs. **h-j**, Dorsal view of confocal stacks of electroporated somites observed at E5.5 (as in a-d). **k**, Column graph for h-j showing the percentage of fusion events relative to controls distributed in myocytes containing the indicated number of nuclei. Striped colour columns indicate non-significant differences with controls. Embryos in a-e and h-j were stained against GFP and RFP antibodies. Arrowheads point to cell nuclei in selected fibres. ***P<0.0001; ** P<0.01 Scale bars: 50μm.

### A TGFβ receptor complementation mechanism triggers a fusion-refractory state.

The cellular mechanism(s) whereby TGFβ signalling may regulate fusion in the embryo is suggested by the expression patterns of TGFβ family members in early somites (Fig. 1f-i) and the known fusion behaviour of different populations of cells. During early myogenesis DML-derived myofibres fuse only to cells emanating from the anterior and posterior borders of the dermomyotome^9^ (Supplementary Fig. 3). Thus, the fusion of (TGFBR2-positive) progenitors emanating from the anterior and posterior borders of the dermomyotome, onto (TGFB2/3- and TGFBR1-positive) myocytes derived from the DML, should, through a receptor complementation process, bring all TGFβ pathway components together, thereby allowing the activation of its signalling, and resulting in a decrease/arrest of fusion.

To test this appealing hypothesis, we first evaluated whether TGFBR1 is regulating the fusion of DML-derived myocytes to progenitors originating from the anterior and posterior borders of the dermomyotome. The expression of MLC-driven constitutively active (CA) and dominant negative (DN) forms of TGFBR1 in DML-derived myocytes resulted in an overall significant reduction or increase, respectively, in their fusion when compared to controls (*** P-value<0.001; Fig. 3a-e). We observed for CA TGFBR1 an increase (+22%, compared to controls) in mononucleated fibres and a decrease (−72%, compared to controls) in myocytes containing 4 and more nuclei (Fig. 3c, e). In contrast, DN TGFBR1 resulted in a decrease (−34%, compared to controls) in mononucleated myocytes and an increase (+95%, compared to controls) in large fibres (Fig. 3d-e). Similar to DN-TGFBR1, a CRISPR/Cas9-mediated loss-of-function against TGFBR1 significantly increased fusion (*** P-value<0.001; Fig. 3f, g, i), resulting in a decrease (−49%, compared to controls) in mononucleated fibres and an increase (+98%, compared to controls) in large fibres. These experiments demonstrate that TGFBR1 is the type I receptor regulating fusion in somites. TGFBR1 heterodimerizes with TGFBR2 to transmit signals from TGFβ ligands intracellularly^11^. If TGFBR2 expressed by anterior and posterior border-derived cells were responsible for triggering the TGFβ pathway and inhibiting fusion in DML-derived fibres, its loss-of-function in anterior or posterior border-derived cells should lead to an increase of fusion, while its loss-of-function in DML-derived cells should have no effect. Similarly to the DML cell population, the anterior or posterior borders can be specifically targeted by electroporation^19^. We first electroporated the DML with a CRISPR/Cas9 construct against TGFBR2; quantification of fusion showed that this did not significantly modify fusion when compared to controls (P-value: 0.89; Fig. 3h-i). In contrast, electroporation of the same CRISPR/Cas9 TGFBR2 construct in the posterior border showed a significant increase in overall fusion of myocytes (*** P-value<0.001) with a decrease (−13%, compared to controls) in mononucleated fibres and an increase (+46%, compared to controls) in large myocytes containing 4 or more nuclei (Fig. 3j-l). Based on this data, we propose a model whereby the migration of anterior or posterior border myoblasts and their fusion to DML-derived myocytes results in the activation of the TGFβ pathway by a receptor heterodimer complementation mechanism, ultimately leading to the inhibition of fusion.

**Figure 3.**
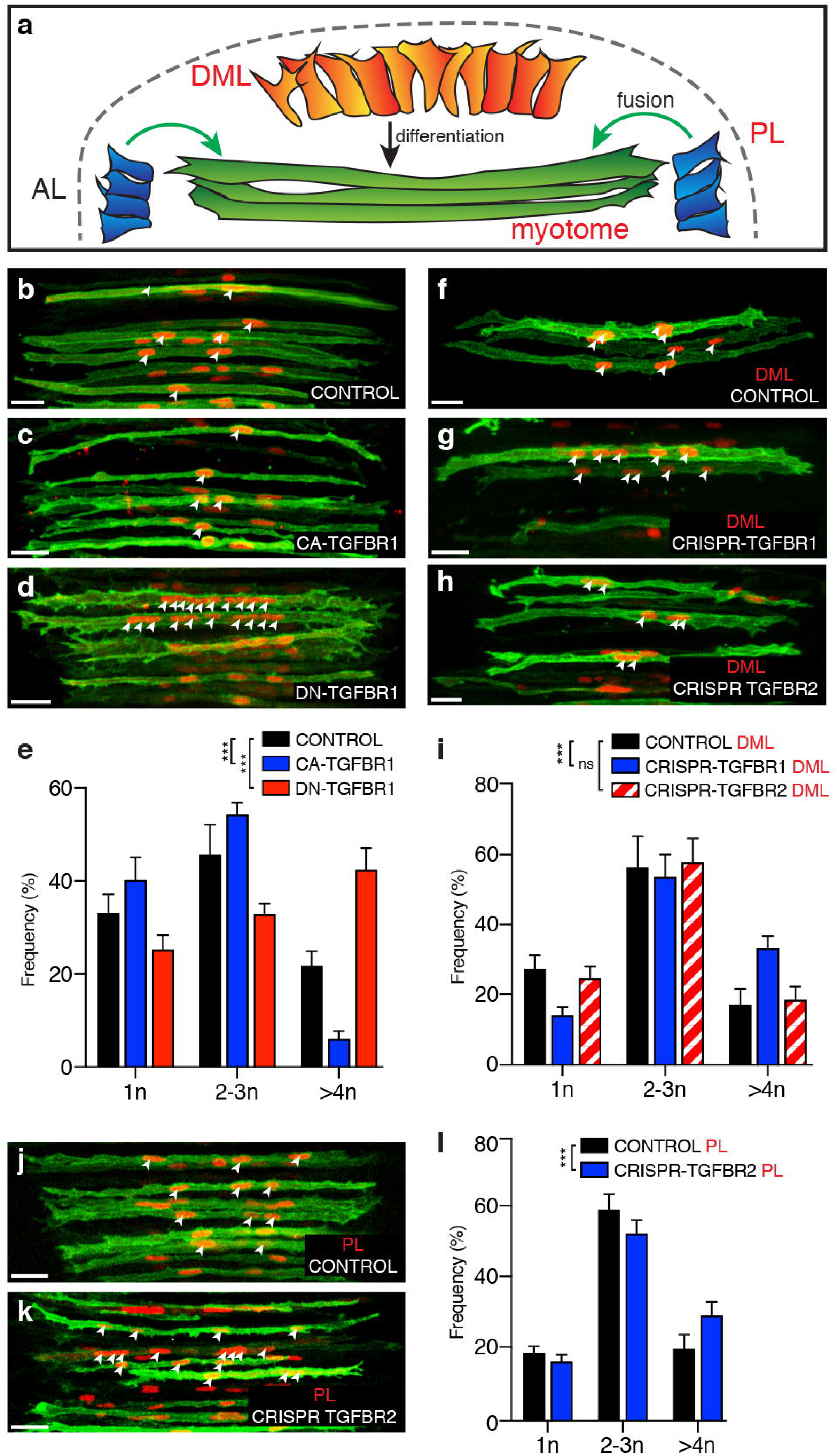
TGFBR1 in DML-derived myocytes and TGFBR2 in posterior dermomyotome border progenitors are necessary for fusion. **a**, schematics showing the different structures targeted in these experiments. The DML generates the myocytes of the myotome, the anterior (AL) and posterior (PL) borders provide progenitors that fuse to DML-derived myocytes **b-d**, DML-derived myocytes electroporated with MLC promoter driving expression of constitutively active (CA, c) and dominant-negative (DN, d) TGFBR1 variants, together with membrane GFP (green) and nuclear mCherry (red). **e**, Column graph for b-d showing the population of electroporated myocytes containing the indicated number of nuclei relative to their controls (in %). **f-h**, DML-derived myocytes electroporated with a CRISPR/Cas9 construct targeting the chicken TGFBR1 or TGFBR2 sequences together with membrane GFP (green) and nuclear mCherry (red). **i**, Column graph for f-h showing the population of electroporated myocytes containing the indicated number of nuclei relative to their controls (in %). Solid colour columns indicate overall significant difference with controls. Striped colour columns indicate non-significant difference with controls. **j,k**, Myocytes electroporated in the posterior border of the dermomyotome with a CRISPR/Cas9 construct targeting the chicken TGFBR2 sequence together with membrane GFP (green) and nuclear mCherry (red). **l**, Column graph for j-k showing the population of electroporated myocytes containing the indicated number of nuclei relative to their controls (in %). Embryos in b-d, f-h and j-k were fixed at E5.5 and stained against GFP and RFP antibodies. Arrowheads point to cell nuclei in select fibres. ***P<0.0001, ns = p>0.05. Scale bars: 50μm.

### Fusion-competence is restored through is RAB-dependent endocytic recycling of receptors

Since all myofibres within the myotome eventually become polynucleated, we reasoned that an inhibition of fusion during development can only be temporary. Therefore, to switch from a fusion-refractory state to a fusion-competent state, it is crucial that TGFβ signalling decreases within the receiving myofibre, thereby allowing for additional rounds of myoblast fusion to occur. Thus, we asked whether myoblast fusion-refractory and fusion-competent states could be controlled by cycling TGFβ receptor activity. Like other cell surface receptors, TGFβ receptors are rapidly internalized after activation. The main mechanism of TGFBR internalization is via clathrin-dependent endocytosis^30^. The small GTPase RAB proteins are key regulators of intracellular membrane trafficking of growth factor receptors^31^. Both type I and type II TGFβ receptors are recycled back to the cell surface by recycling endosomes, through a RAB11-dependent mechanism. On the contrary, lysosomal degradation, mediated by a RAB7-dependent mechanism, terminates signal transduction. Inhibiting RAB11 function should thus lower receptor recycling at the cell membrane, and ultimately decrease receptor signalling. Conversely, inhibiting RAB7 function should reduce receptor degradation, resulting in an extension of signal transduction. To test this, we electroporated a dominant-negative form of RAB11 (under the MLC promoter) in DML-derived fibres. Three days later, this resulted in a significant increase (*** P-value<0.001) in fusion when compared to controls (Fig. 4a-c), resulting in a decrease (−34%, compared to controls) in mononucleated fibres and an increase (+203%, compared to controls) in large fibres. In contrast, electroporation of DN-RAB7 in DML-derived fibres led to a significant reduction (*** P-value<0.001) in fusion when compared to controls (Fig. 4d-f), resulting in more (+59%, compared to controls) mononucleated fibres and less (−51%, compared to controls) in large myocytes. Altogether these results suggest that in muscle fibres, the recycling or conversely, the degradation of TGFβ receptors play important roles in the regulation of myoblast fusion to myofibres. Furthermore, this supports the hypothesis that consecutive fusion-refractory and fusion-competent states are controlled by cyclic TGFβ signalling, effectively regulating the pace of myoblast fusion in vertebrate embryos.

**Figure 4.**
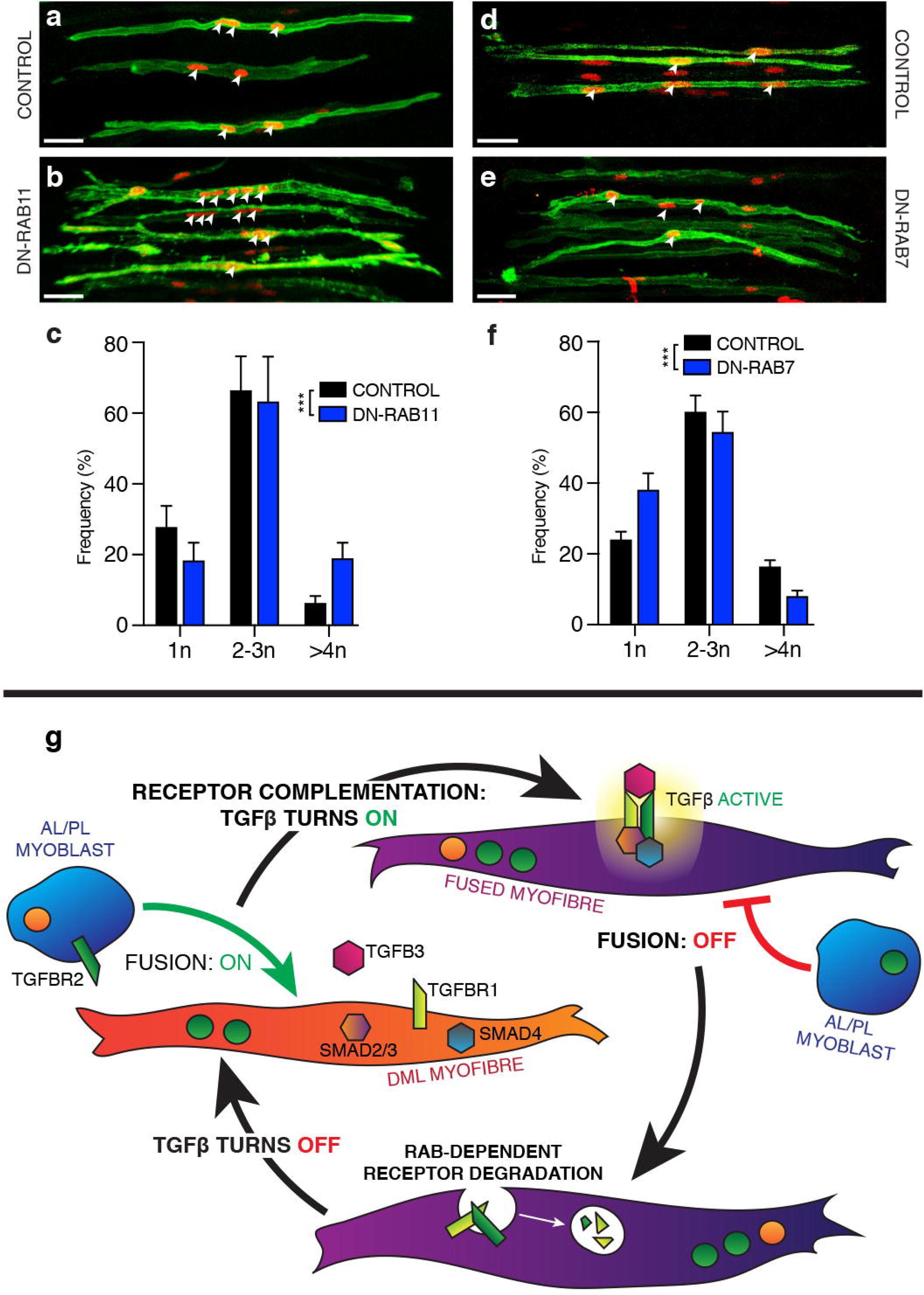
Myoblast fusion is dependent upon RAB-dependent endocytic recycling of receptors. **a-b, d-e**, DML-derived myocytes electroporated with MLC promoter driving expression of dominant negative (DN) variants of RAB11 (b) and RAB7 (e), together with membrane GFP (green) and nuclear mCherry (red). **c,f**, Column graph for a-b and d-e showing the population of electroporated myocytes containing the indicated number of nuclei relative to their controls (in %). **g**, A model describing the receptor complementation mechanism: prior to fusion, myocytes originating from the DML (in orange) express the ligand TGFB3, the effectors SMAD 3, & 4 and the receptor TGFBR1. They are competent to fuse. Myoblasts originating from the posterior or anterior borders of the somite (in blue) express TGFBR2. Upon fusion of myoblasts to DML-derived myocytes, membrane merging of the fusion partners allow the activation of the TGFβ pathway by a receptor heterodimer complementation mechanism (in purple), ultimately leading to a temporary inhibition of fusion of another myoblast. Return to a fusion-competent state is dependent upon RAB-mediated receptor degradation.***P<0.0001. Scale bars: 50μm.

The present study demonstrates (Fig. 4g) that TGFβ signalling plays a major role in the morphogenesis of skeletal muscles during embryogenesis. In contrast to the previously identified genes belonging to the machinery necessary for fusion, the TGFβ pathway acts as a molecular brake in this process. Furthermore, the hyper-fusion phenotype that we obtained when releasing its signalling and the observation that many tested genes inhibited the fusion of C2C12 *in vitro* suggests that tight harnessing of fusion at multiple molecular check-points is an unsuspectedly crucial aspect of muscle formation. The demonstration that unregulated fusion is detrimental to muscle function in rodents and that the TGFβ pathway plays an identical function in adult mice (Girardi et al. BiorXiv, 2019) supports the hypothesis that the inhibitory function of TGFβ signalling on myoblast fusion is critical to many if not all aspects of muscle growth and repair in vertebrates.

The technology of *in vivo* electroporation in the chicken embryo allows an unmatched control of the timing and location of gene function, thus providing a powerful approach to characterize the mechanisms that coordinate cell signalling with cell and tissue movements in developing amniotes. The mechanism we uncovered here is unique as it relies on the fusion of progenitors to myofibres, which allows receptor dimer complementation and signalling to occur. The resulting inhibition of fusion suggests that auto-inhibition is the mechanism whereby fusion controls itself. The reversible nature of receptor signalling (through RAB-mediated receptor degradation) completes this mechanism, allowing it to reset. Therefore, membrane fusion, signalling and receptor degradation collaborate in a reiterating regulatory module that ultimately controls whether fusion takes place and at what pace.

Altogether these findings uncover that muscle progenitors, progressing through the myogenic program and thereby acquiring fusion competency, are confronted with an additional level of regulation, epitomized by TGFβ signalling. Acting as a molecular brake, it expands the possibilities of temporal and spatial control of muscle patterning, from a simple stop or go signal to the fine-tuning of fusion that likely governs crucial phases of muscle growth and regeneration. This opens new routes of investigation into an unforeseen aspect of muscle biology that could bring important benefits to muscle therapies.

## Acknowledgments

We thank Monash Micro Imaging (MMI) and the Centre d’Imagerie Quantitative Lyon-Est (CIQLE) for imaging support; Profs. Peter Currie and Olivier Pourquié for critical reading of the manuscript. This work was funded by grants from the National Health and Medical Research Council (NHMRC, Australia), by the Australian Research Council (ARC, Australia), by the Association Française contre les Myopathies (AFM, France) and by the EU 6th Framework Programme Network of Excellence MYORES. The Australian Regenerative Medicine Institute is supported by grants from the State Government of Victoria and the Australian Government.

## Author Contributions

CM, DSieiro and JM conceived and designed the study. DSieiro, JM and VM performed all experiments, DSalgado performed all the bioinformatic treatment of data. CM and DSieiro wrote the manuscript. All authors reviewed the manuscript.

## Footnote

Present addresses: Daniel Sieiro, Department of Pathology Brigham and Women’s Hospital Harvard Medical School, Boston, MA, USA. David Salgado: Aix Marseille University, Faculty of Medicine La Timone, INSERM U1251, Marseille, France.

## Supplementary Figure legends

**Supplementary Figure 1**

**Representation of the predicted protein-protein interactions between candidate genes inhibitory (in red) and necessary (in green) for C2C12 myoblast fusion.** The 200 most significant genes for each category were analysed through String software^32^, using highest confidence interaction score. Disconnected nodes were hidden from final graph. Fusion-inhibitory and -necessary molecules were pseudo-coloured manually using Adobe Illustrator. Statistical overrepresentation tests (PANTHER and INGENUITY) on Gene Ontology molecular functions identified the TGFβ family as overrepresented (circled).

**Supplementary Figure 2**

**Diagram depicting essential molecules involved in canonical TGF**β **signalling.** TGFβ ligands (BMPs, GDFs, TGFβs, among others) bind type 2 receptors, which in turn recruit a type 1 receptor to form a hetero-tetrameric complex. Ligand-bound receptor 2 phosphorylates the serine residues of the receptor 1, which in turn phosphorylates the effector proteins R-SMADs (Receptor-regulated SMADs). BMP receptors only activate SMADs 1/5/8, while TGFBs, Activins and GDFs act through SMADs 2/3. Phosphorylated R-SMADs have a high affinity to SMAD4. The SMAD complex enters the nucleus and binds co-factors and transcription promoters to induce DNA transcription. I-SMADs (Inhibitory SMADs) are involved in negative feedback. SMAD7 competes for receptor 1 binding sites and prevents phosphorylation. SMAD6 competes with SMAD4 exclusively through the BMP SMAD1/5/8 interactions. Diagram has been simplified for clarity and relevance.

**Supplementary Figure 3**

**Diagram depicting the somitic regions taking part in the formation of the myotome and the fusion of myocytes.** Selected epithelial cells originating from the medial border of the dermomyotome (DML) undergo an epithelial to mesenchyme transition (EMT) that allows their translocation in a region located beneath the dermomyotome, the transition zone (TZ), where they orient in the antero-posterior axis of the embryo^19–21^. The EMT triggers the entry of cells derived from the DML into the myogenic program. Terminal myogenic differentiation (e.g. MyHC expression) is observed when cells attach to the anterior and posterior borders of somites, at which time they are named myocytes (green elongated fibres). Fusion of myocytes is observed about 24 hours after myotome formation was initiated. Progenitors from the anterior (AL) and posterior (PL) border of the dermomyotome translocate in the myotome where they fuse to existing myocytes^9^. nt: neural tube.

**Supplementary Figure 4**

**Myosin Light Chain promoter is expressed in terminally differentiated myocytes. a**, Overlay confocal stacks of a single somite electroporated at HH15 (E2.5), and imaged at HH25 (E4.5). The white solid line delineates the somite. The dotted lines delineate the transition zone (TZ). **b**, Electroporation control, a CAGGS ubiquitous promoter driving nuclear H2B-RFP. Expression is observed in the dorsomedial lip (DML), the TZ and the myotome. **c**, Expression of GFP driven by a myosin light chain (MLC) promoter. No expression of the reporter is seen in the DML. Cells within the DML initiate MYF5 and MYOD expression^21^ before entering the TZ. TZ cells are all postmitotic and express MyoG (our observation). Faint GFP expression was observed in a few cells of the TZ, while all myocytes of the myotome robustly expressed the fluorescent marker. MLC is therefore activated in terminally differentiating progenitors; **d**, Antibody staining against Myosin Heavy Chain (MyHC) showing expression exclusively in differentiated muscle fibres of the myotome. RFP and GFP were detected with IHC. Scale bars: 50μm.

**Supplementary Figure 5**

**Activation or inhibition of TGF**β **signalling does not affect terminal myogenic differentiation.**

Confocal stacks of somites 3 days after electroporation of SMAD3, SMAD4 (r-Smads effectors) and SMAD7 (i-Smad inhibitor), which severely affect fusion (decrease: SMAD3 and 4 or increase: SMAD7). In all experiments performed (not all shown here), MyHC staining was done in order to ensure terminal differentiation was not being affected. Regardless of fusion, MyHC expression was never affected. White arrows point to select electroporated fibres demonstrating a normal expression of MyHC. Scale bars: 50μm.

**Table 1**

Scores (as normalized to 100 to the effect of negative controls, RLuc and FLuc), accompanied with their respective P-values, obtained after esiRNA transfection of the indicated genes into the mouse myogenic cell line C2C12. Effects on fusion (nuclei per fibre, in red), proliferation/cell survival (nuclei count, in blue) and myogenic differentiation (myogenin expression, in green) after loss of function of those molecules are indicated. The list encompasses the gene names (Entrez gene IDs) of the esiRNAs that obtained the highest score in the fusion assay. The corresponding genes are therefore predicted to act as strongest inhibitors of C2C12 fusion.

**Table 2**

Scores (as normalized to 100 to the effect of negative controls, RLuc and FLuc), accompanied with their respective P-values, obtained after esiRNA transfection of the indicated genes into the mouse myogenic cell line C2C12. Effects on fusion (nuclei per fibre, in red), proliferation/cell survival (nuclei count, in blue) and myogenic differentiation (myogenin expression, in green) after loss of function of those molecules are indicated. The list encompasses the Entrez gene IDs of the esiRNAs that obtained the lowest score in the fusion assay. The corresponding genes are therefore predicted to act as most necessary for C2C12 fusion.

**Table 3**

List of tested members of the (SMAD2/3 dependent) TGFβ signalling pathway (TGFBs, TGFBRs, SMADs, including activins and inhibins) of molecules known to synergize with this pathway^33–36^ and of BMP (SMAD1/5/8) related genes. Score calculation as described in tables 1 and 2.

## Methods

### EsiRNA screen generation and candidate selection

We performed a genome-wide RNA interference (RNAi) functional screen on a mouse C2C12 muscle cell line (Fig. 1). The strategy was based on the transfection of long dsRNA enzymatically digested by bacterial RNase III for the generation of “esiRNA”. This process generates a heterogeneous population of siRNAs capable of interacting with multiple sites on the target mRNA: since each single siRNA in a pool has different off-targets while having the same on-target, the use of siRNA pools dilutes out off-target effects^37–39^. The esiRNA library comprised 8628 mouse genes. An EGFP-positive C2C12 cell line was transfected with esiRNAs, and they were placed in differentiation medium. Three days later, the detection of fused myotubes was performed, using an automated cell-recognition imaging program (Definiens) that quantified the number of nuclei (stained with Hoechst) per fibre (detected by the EGFP green fluorescence). Since a lack of C2C12 fusion upon esiRNA transfection could be a true effect on cell fusion or due to a failure to differentiate, the wells were also stained for myogenin to evaluate the myogenic differentiation at the end of the experiment. The total number of nuclei per well also gave an indication that the esiRNAs did not interfere with cell proliferation or survival. In each plate, a number of control wells served to normalize the results obtained in each assay. They were transfected with esiRNA against Renilla and Firefly Luciferase (RLuc and FLuc, negative controls) and against the Rac Family Small GTPase 1 and the Cell Division Cycle 42 genes (RAC1 and CDC42, positive controls). We have performed two rounds of transfection and each esiRNA was tested in triplicate. The readings were normalized (to 100) to the effect of negative control esiRNAs (RLuc and FLuc) assayed on each multi-well plate. As expected, transfection of RAC1 and CDC42 esiRNAs led to a significant decrease in the fusion of C2C12 (RAC1: −27%; CDC42: −48% compared to controls, Table 3).

### In situ hybridization and sectioning

*Insitu* hybridizations were performed as described in (^20^). RNA probes were generated from ESTs obtained from the chick EST database at BBSRC^40^. The following EST was used for TGFBR2: chEST327g19. Images for *insitu* hybridizations of TGFB2, TGFB3 and TGFBR1 were obtained with permission from the GEISHA database^41^. Embryos were either analysed after whole-mount staining or sectioned into 15μm slices using a Leica cryostat (as described in^42^).

### Chicken electroporation

The somite electroporation technique that was used throughout this study has been described previously (^21,43^). Chicken embryos at stage 15–16 of Hamburger Hamilton (HH^17^; 24–28 somites) were electroporated in the dorso-medial border (DML) or the posterior border (PL) of newly formed interlimb somites. The electroporation technique results in the mosaic expression (50% at best^9^) of the DNA constructs in the somite sub-domains (DML and PL). This is particularly important in the case of a study on muscle cell fusion, since the observed phenotypes result from the competition of electroporated and wild-type myoblasts for the fusion to recipient myofibres. This “diluting effect” is even more pronounced in the case of PL electroporation, since PL-derived electroporated myoblasts compete for the fusion to DML-derived myocytes not only with WT myocytes from the PL, but also from WT myoblasts from the AL.

### Expression constructs

The CAGGS-transposase and the Tol2 MLC-nls-mCherry were described before^9^. The following plasmids were created by inserting in frame each sequence described below into a Tol2 MLC-IRES GFPcaax (i.e. an EGFP addressed to the plasma membrane by a CAAX-box prenylation domain). The mouse SMAD43, SMAD4, SMAD6 and SMAD7 were obtained from the I.M.A.G.E. consortium (distributed by Source BioScience). A mouse SMAD5 was obtained from pCMV5-Smad5 (gift from Jeff Wrana, Addgene plasmid #11744^44^). A constitutively active form (CA SMAD5) was generated by replacing Serine 463 and 465 by Aspartic Acid (S463D, S465D). A constitutively active form of the human SMAD1 (S463E, S465E) was obtained from pCS2 hSmad1-EVE (gift from Edward De Robertis, Addgene plasmid #22993^45^). A constitutively active form of the rat TGFBR1 (T202D) was obtained from pRK5 TGF beta type I receptor (gift from Rik Derynck, Addgene plasmid #14833^46^). A dominant-negative form of the human TGFBR1 (K232R) was obtained from pCMV5B-TGFbeta receptor I K232R (gift from Jeff Wrana, Addgene plasmid #11763^47^). A dominant negative form of the human Rab11A (S25N) was obtained from plasmid PCMV-intron myc Rab11 S25N (a gift from Terry Hébert, Addgene plasmid #46786^48^). A dominant negative form of the human Rab7A (T22N) was obtained from plasmid DsRed-rab7 DN (a gift from Richard Pagano, Addgene plasmid #12662^49^).

### CRISPR/Cas9 constructs

The use of CRISPR/Cas9 technology on somatic cells in chicken was based on (^25,26^). Guide RNA design was done using the site CHOPCHOP: http://chopchop.cbu.uib.no/ (Cornell, Montague^50^). The gRNA sequences were as follow: for the chicken SMAD3: gRNA1 cSMAD3: gTCATCTACTGCCGGCTGTGG (targets exon 2); gRNA2 cSMAD3: gTCCACTCGTTGGTAATGATA (targets exon 2); for the chicken TGFBR1: gRNA1 cTGFBR1: caccGTAACTCCATCTCTGAAGGA (targets exon 2); gRNA2 cTGFBRR1: caccgAGTATTGGTAAGGGTCGCTT (targets exon 4); for the chicken TGFBR2: gRNA1 cTGFBR2: caccGCTTCCTCTTCTTGTGAGTG (targets exon 4); gRNA2 cTGFBR2: caccGTGGATGACTTGGCCAACAG (targets exon 4). Two sites per gene were chosen (hence the two gRNA sites) in order to excise a portion of the coding sequence and promote the loss of function of the targeted proteins in electroporated cells. The expected genomic DNA deletions sizes are: cSMAD3: 136bp; cTGFBR1: 8096bp; cTGFBR2: 681bp. The efficiency of the chosen gRNAs was tested by transfection of the constructs in DF1 chicken fibroblast cells *in vitro* (as described in ^25,26^).

### Immunohistochemistry and imaging

Whole-mount antibody staining were performed as described (^20^). The following antibodies were used: rabbit polyclonal directed against RFP (Abcam #62341, 1/1000), chicken polyclonal antibody against EGFP (Abcam #13970, 1/1000) and mouse monoclonal directed against Myosin Heavy Chain MYH1 (MF20) (deposited to the DSHB by Fischman, D.A.). Embryos were washed and cleared in 80% glycerol/PBS. Whole-mount embryos were imaged using a Leica SP5 confocal microscope with scanning resonance running LAS AF software (Leica MicroSystems). Image stacks were analysed by using either the Imaris software package (Bitplane, version 7.5.2) or ImageJ software.

### Quantification and statistics

Manual counting of nuclei per myocytes within confocal z-stacks was performed using the cell counter plugin of ImageJ (^51^). Statistical analyses were performed using the GraphPad Prism software. Each treatment comprised about 6 embryos from two independent experiments, and about 3 somites per embryo. The number of counted myocytes per experimental condition was about 300 fibres (6150 fibres total). Mann-Whitney non-parametric tests were applied on the entire population of counted myofibres to evaluate significance of each treatment (P-values are indicated in the text). Experimental treatments (gain or loss of gene function) that led to significant differences compared to controls are indicated on graphs by solid colour columns; non-significant results are indicated by striped colour columns; controls are indicated by black columns. The column graphs in Fig. 1-4 depict the population of electroporated myocytes that were calculated for each experimental condition. They were distributed in three groups, myocytes that contained one nucleus, those that contained 2-3 nuclei and those containing 4 or more nuclei, expressed as a percentage of the entire myocyte population. Tendencies to decrease or promote fusion are more pronounced in the first and last (1N and >4N) categories. Therefore, we describe in the text the changes, compared to controls, in the proportion of fibres in these two categories. Columns are accompanied by the standard error of the mean (SEM). *** P-value<0.001 extremely significant; ** P-value 0.001 to 0.01 very significant; P-value>0.05 non significant.

