## Supplementary figures and images for "Auto-inhibition of myoblast fusion by cyclic receptor signalling"

### Supplementary Figure 1

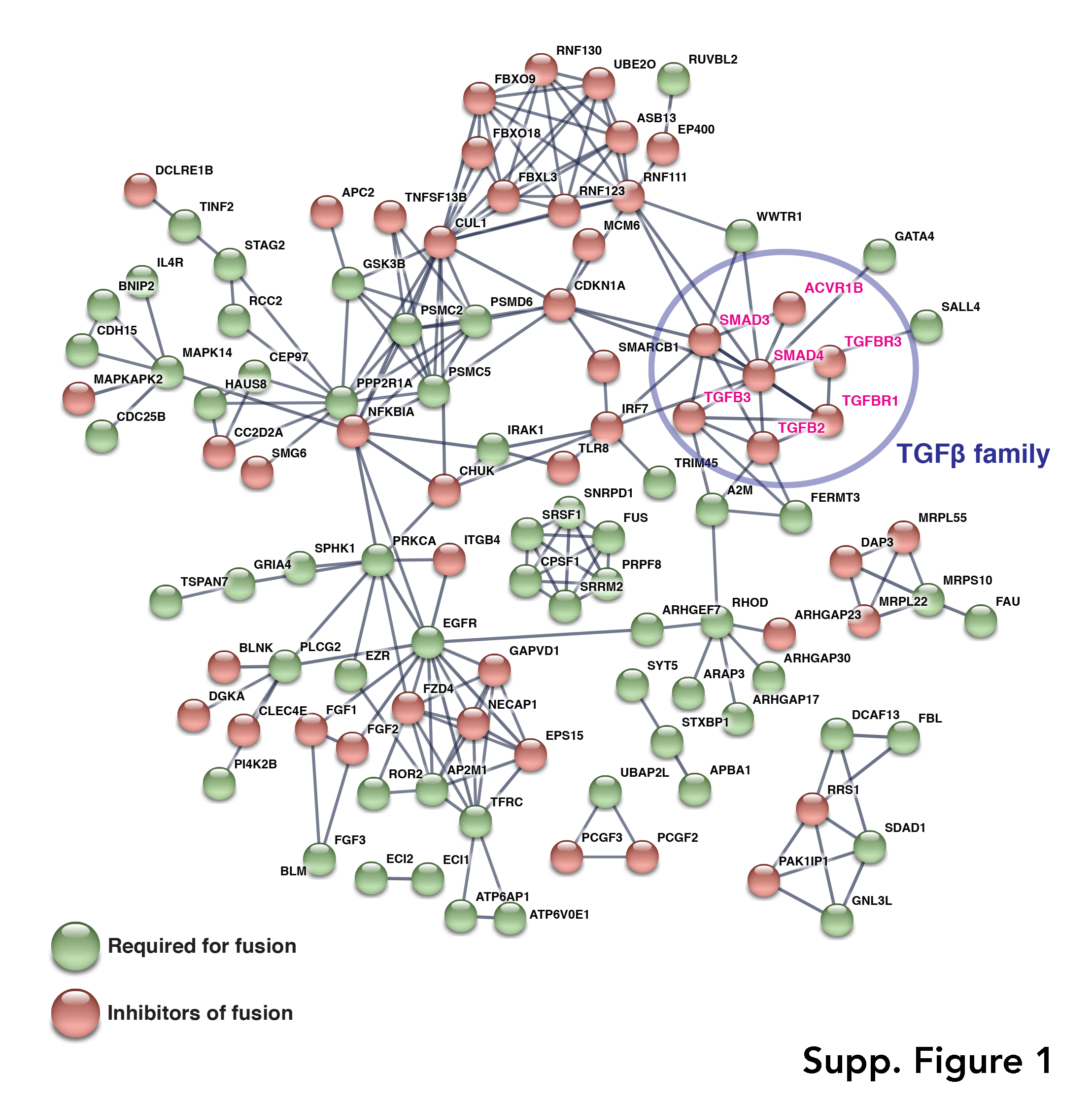

### Supplementary Figure 2

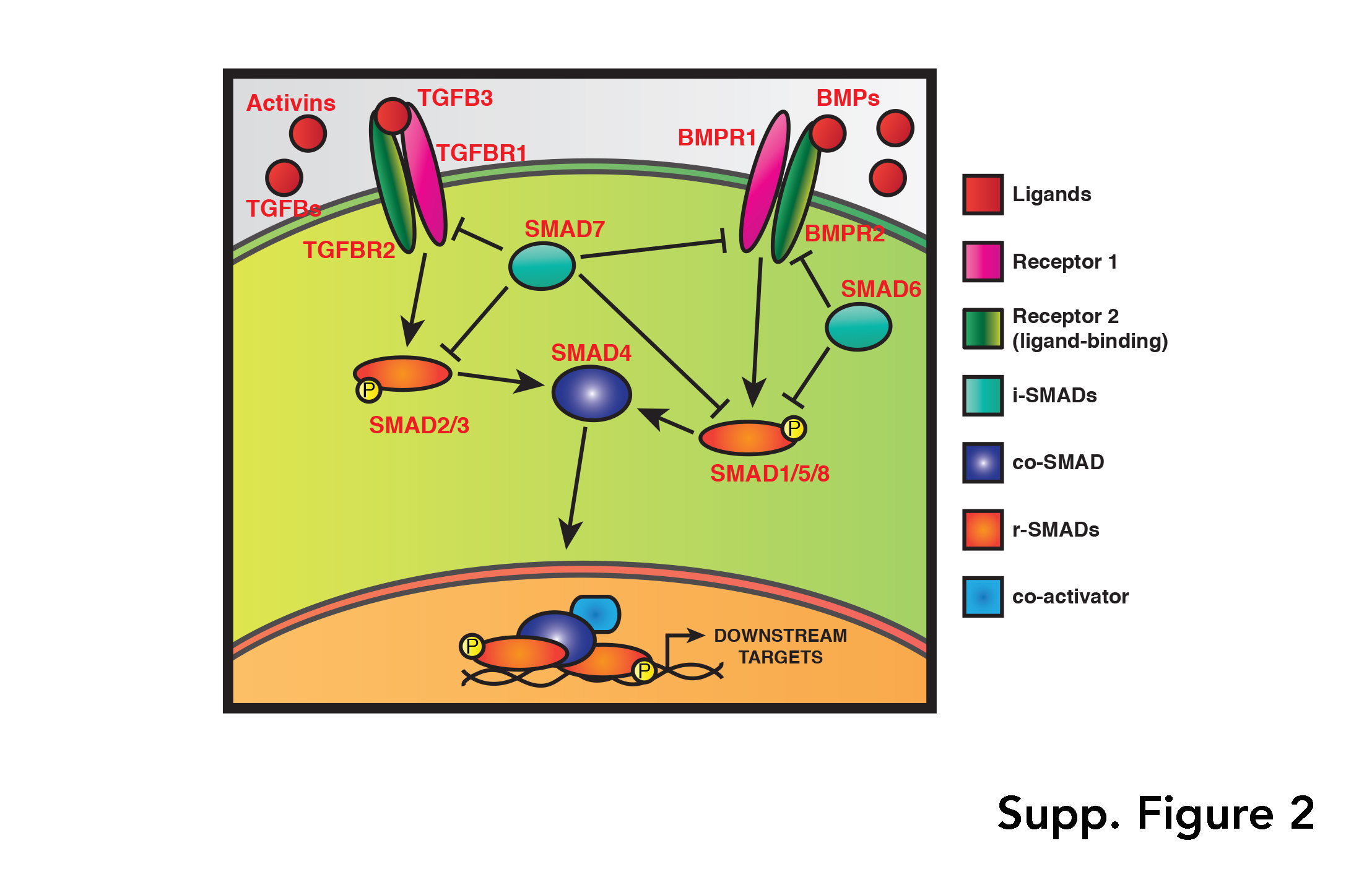

### Supplementary Figure 3

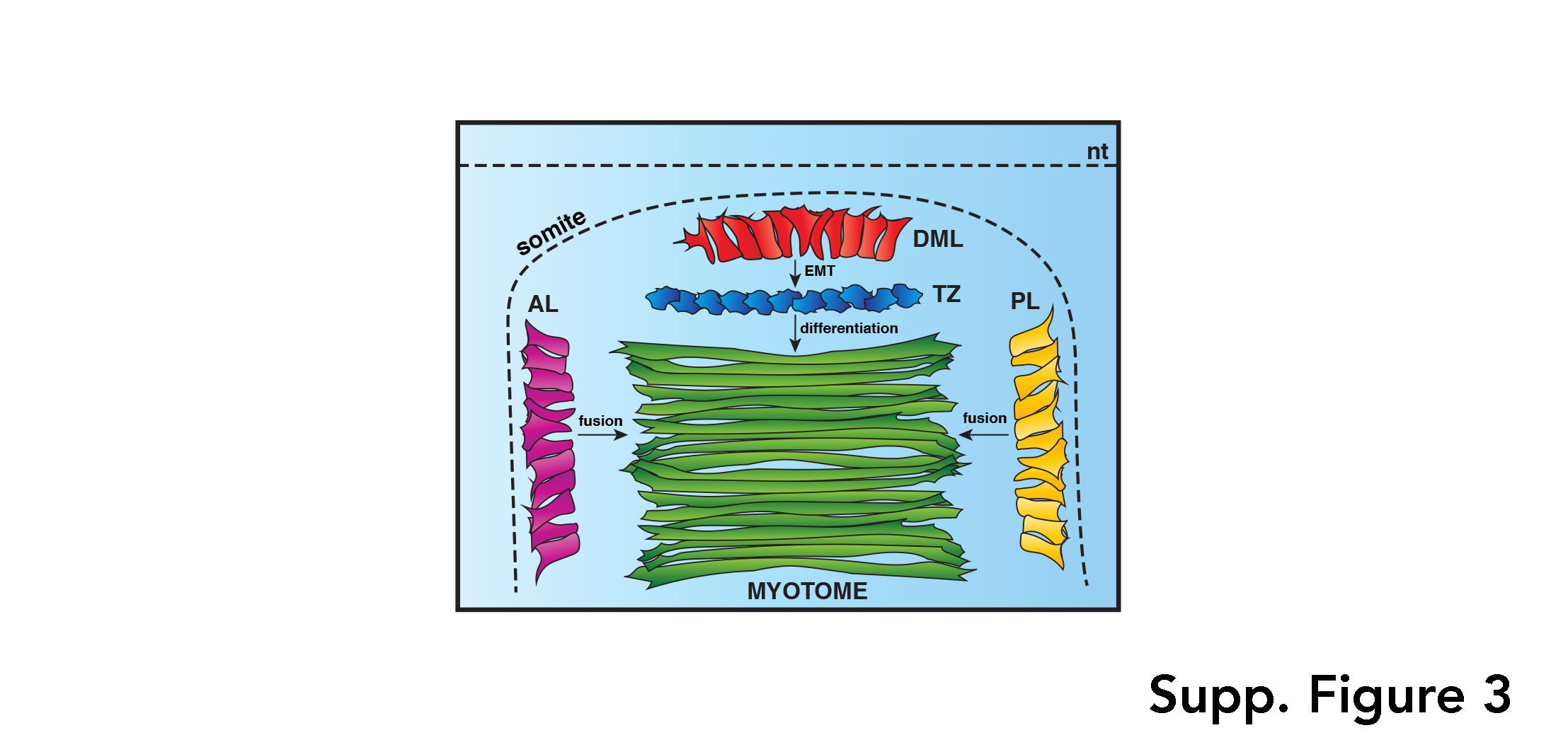

### Supplementary Figure 4

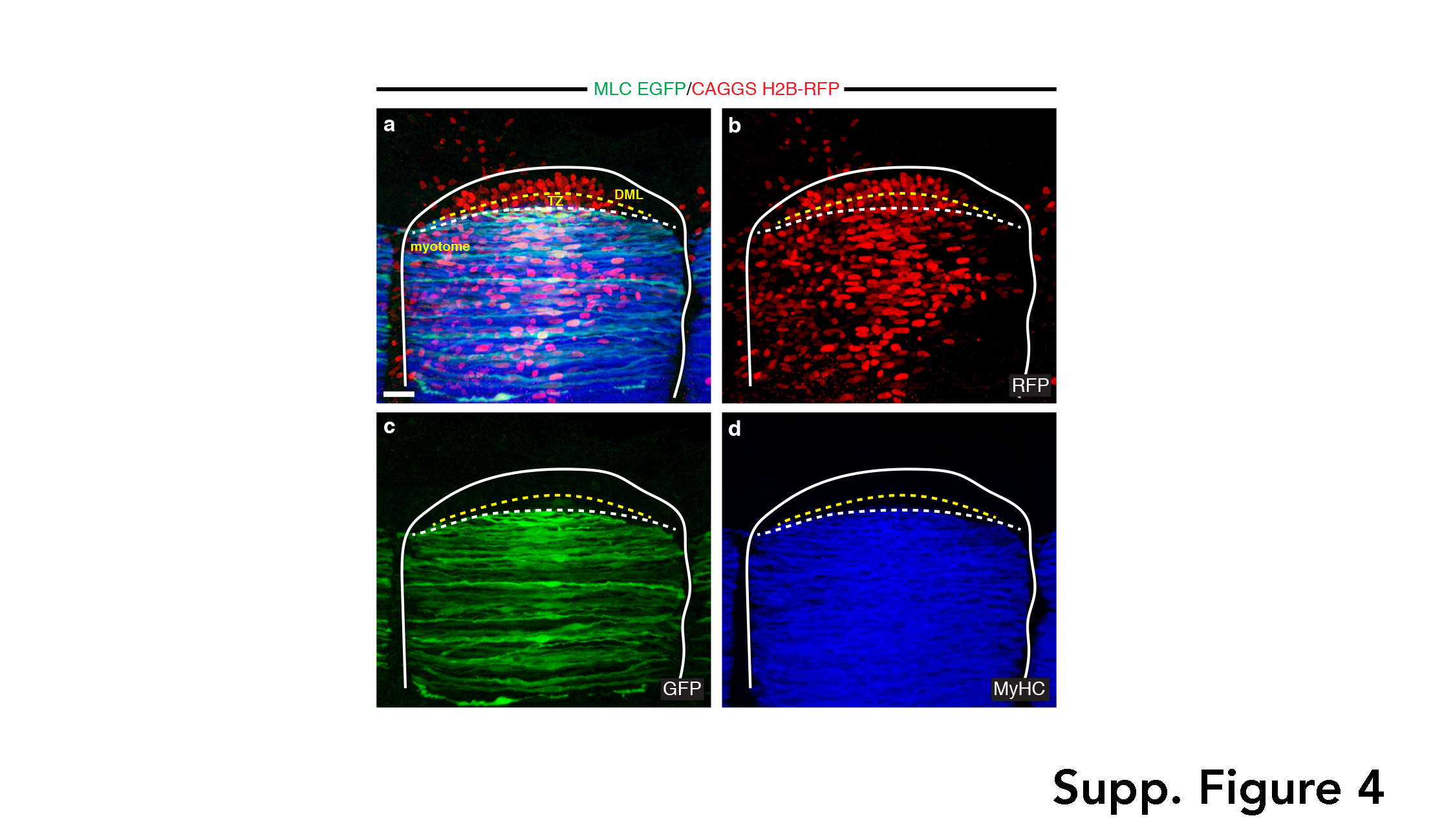
