## Supplementary material for "Auto-inhibition of myoblast fusion by cyclic receptor signalling": Table 1

### Table 1. Genes inhibitors of C2C12 fusion

| Entrez Gene ID | pvalue | %nuclei/myotube | pvalue | %nuclei | pvalue | %myogenin |
|---|---|---|---|---|---|---|
| Smad4 | 8.42E-15 | 525.54 | 0.14 | 98.56 | 2.20E-16 | 156.57 |
| Fzd4 | 2.20E-16 | 508.18 | 0.00 | 111.16 | 2.20E-16 | 139.43 |
| Mdfi | 2.20E-16 | 432.86 | 7.11E-15 | 148.72 | 6.41E-10 | 114.60 |
| Tgfb2 | 6.13E-13 | 405.95 | 0.00 | 111.81 | 2.20E-16 | 154.19 |
| Mmp17 | 2.52E-13 | 400.40 | 0.00 | 115.77 | 2.20E-16 | 145.97 |
| Adh5 | 2.20E-16 | 387.35 | 1.21E-09 | 126.46 | 2.20E-16 | 125.79 |
| Sgcd | 3.77E-11 | 386.64 | 4.84E-07 | 119.80 | 0.01 | 106.67 |
| Cyfip2 | 1.53E-05 | 382.66 | 0.00 | 92.34 | 0.00 | 104.30 |
| 2310002L13Rik | 1.20E-15 | 379.80 | 2.79E-12 | 129.66 | 2.90E-09 | 115.36 |
| Smad3 | 2.20E-16 | 377.95 | 0.73 | 100.32 | 2.20E-16 | 136.95 |
| Habp4 | 1.84E-14 | 375.01 | 2.05E-07 | 85.37 | 2.20E-16 | 152.15 |
| Mrpl55 | 1.66E-12 | 372.03 | 0.00 | 123.44 | 1.05E-13 | 132.64 |
| Pard6b | 2.20E-16 | 360.21 | 2.20E-16 | 145.19 | 0.08 | 104.93 |
| Inpp4a | 4.51E-09 | 348.96 | 1.96E-07 | 121.85 | 3.89E-13 | 130.06 |
| D17Wsu104e | 7.58E-13 | 348.92 | 2.80E-06 | 112.08 | 2.20E-16 | 138.39 |
| Ube2o | 9.79E-13 | 340.50 | 0.22 | 104.49 | 6.23E-08 | 83.31 |
| Srp54a | 8.88E-12 | 334.81 | 2.57E-12 | 119.38 | 1.74E-06 | 111.15 |
| Spaca1 | 1.74E-14 | 329.89 | 5.73E-12 | 131.26 | 6.11E-16 | 136.73 |
| Best1 | 2.20E-16 | 324.30 | 6.24E-05 | 110.93 | 0.08 | 93.83 |
| Rnf111 | 8.86E-16 | 319.44 | 0.25 | 103.26 | 2.20E-16 | 136.57 |
| 3110062M04Rik | 2.85E-14 | 307.52 | 1.87E-05 | 113.99 | 2.72E-16 | 130.69 |
| Dnaic1 | 2.96E-15 | 304.79 | 1.16E-09 | 121.40 | 2.20E-16 | 135.97 |
| Osbpl8 | 2.20E-16 | 300.28 | 6.92E-06 | 115.59 | 2.20E-16 | 143.16 |
| Qpctl | 1.96E-11 | 295.90 | 8.18E-09 | 127.91 | 3.18E-15 | 128.44 |
| Capn7 | 2.20E-16 | 295.42 | 6.03E-09 | 116.37 | 2.21E-14 | 122.16 |
| 6330416G13Rik | 7.27E-16 | 293.11 | 4.84E-10 | 117.67 | 2.80E-05 | 109.34 |
| Tgfbr1 | 2.06E-13 | 284.01 | 0.56 | 97.57 | 2.20E-16 | 148.79 |
| Rrs1 | 9.95E-12 | 280.82 | 4.41E-07 | 117.55 | 2.20E-16 | 145.62 |
| Fis1 | 4.01E-12 | 280.12 | 1.90E-11 | 132.70 | 1.31E-13 | 72.58 |
| Csrnp1 | 7.00E-13 | 276.67 | 3.40E-14 | 134.42 | 1.85E-11 | 121.61 |
| Ep400 | 2.55E-13 | 276.09 | 0.71 | 99.15 | 4.31E-14 | 125.76 |
| Mpv17l | 1.33E-14 | 275.41 | 0.00 | 110.83 | 2.20E-16 | 148.62 |
| 2410018M08Rik | 4.33E-15 | 274.92 | 0.01 | 107.24 | 2.20E-16 | 130.66 |
| Pcgf2 | 6.21E-13 | 274.88 | 3.39E-11 | 119.28 | 2.20E-16 | 139.64 |
| Tnfsf13b | 1.76E-09 | 271.19 | 0.33 | 102.68 | 3.62E-16 | 125.97 |
| Fam167b | 2.31E-11 | 271.15 | 5.74E-13 | 130.48 | 7.14E-16 | 126.39 |
| Scamp2 | 1.51E-13 | 269.26 | 3.09E-10 | 124.17 | 0.01 | 107.04 |
| Wipi2 | 1.60E-09 | 268.12 | 0.00 | 115.89 | 2.20E-16 | 136.86 |
| Slc9a4 | 1.30E-14 | 266.91 | 0.61 | 101.61 | 2.37E-11 | 118.18 |
| Gja10 | 7.42E-15 | 266.80 | 1.15E-09 | 118.39 | 2.20E-16 | 130.90 |
| Tgds | 3.08E-15 | 266.63 | 6.14E-06 | 113.59 | 2.20E-16 | 151.78 |
| Mixl1 | 3.32E-14 | 266.55 | 6.42E-10 | 120.73 | 2.20E-16 | 131.39 |
| Ahsa1 | 3.65E-11 | 265.95 | 0.12 | 102.81 | 2.20E-16 | 138.02 |
| Apc2 | 4.38E-15 | 263.90 | 1.64E-10 | 120.29 | 4.82E-16 | 126.99 |
| Abca9 | 1.03E-12 | 262.27 | 6.74E-07 | 116.35 | 1.43E-15 | 127.14 |
| Pcmtd2 | 6.76E-12 | 261.94 | 0.03 | 108.37 | 2.20E-16 | 133.32 |
| Rab35 | 1.15E-13 | 258.54 | 2.20E-13 | 123.68 | 2.20E-16 | 141.86 |
| Fam122b | 4.42E-16 | 258.27 | 3.00E-04 | 109.98 | 2.20E-16 | 129.91 |
| Smarcb1 | 3.34E-11 | 254.57 | 0.83 | 101.59 | 2.20E-16 | 140.38 |
| Acvr1b | 2.02E-15 | 254.50 | 8.15E-06 | 111.56 | 2.20E-16 | 124.43 |
| Rundc3a | 3.62E-12 | 253.83 | 6.91E-11 | 118.88 | 2.99E-11 | 116.02 |
| Plekha3 | 2.20E-16 | 253.54 | 3.03E-06 | 113.11 | 4.13E-11 | 115.93 |
| Necap1 | 2.00E-15 | 253.42 | 4.26E-13 | 122.07 | 2.20E-16 | 136.07 |
| Olfr1189 | 1.17E-13 | 253.01 | 3.11E-06 | 114.19 | 2.20E-16 | 123.50 |
| Ctps2 | 2.93E-06 | 252.82 | 0.00 | 115.25 | 2.20E-16 | 141.78 |
| Lrit2 | 3.60E-16 | 252.38 | 0.00 | 112.53 | 3.84E-11 | 118.01 |
| Gzf1 | 3.27E-13 | 252.11 | 7.47E-07 | 115.06 | 2.77E-15 | 124.70 |
| Kcnj6 | 4.26E-12 | 249.45 | 1.85E-14 | 129.84 | 0.00 | 92.23 |
| Asb13 | 6.09E-14 | 248.88 | 7.33E-07 | 113.51 | 8.29E-15 | 127.14 |
| Nr1h5 | 1.94E-15 | 248.85 | 0.00 | 108.47 | 2.20E-16 | 135.59 |
| Pcdhb7 | 3.28E-13 | 247.72 | 0.00 | 109.43 | 6.00E-16 | 129.21 |
| Mcm6 | 3.02E-16 | 247.23 | 3.50E-09 | 118.13 | 2.20E-16 | 137.33 |
| Alg5 | 2.02E-14 | 245.67 | 0.04 | 103.72 | 2.20E-16 | 138.85 |
| Mut | 7.92E-14 | 245.64 | 0.02 | 106.53 | 2.20E-16 | 141.59 |

| | | | | | | |
|---|---|---|---|---|---|---|
| Slc25a12 | 2.47E-13 | 245.47 | 4.34E-05 | 109.73 | 1.00E-15 | 125.34 |
| Cbll1 | 6.92E-14 | 241.74 | 7.38E-05 | 111.77 | 2.43E-11 | 117.37 |
| Ube4b | 2.50E-12 | 240.76 | 0.36 | 97.13 | 0.13 | 103.00 |
| Gdap2 | 1.92E-12 | 240.66 | 0.00 | 113.15 | 1.48E-12 | 121.31 |
| Fgf1 | 8.63E-15 | 239.61 | 4.93E-16 | 128.92 | 2.20E-16 | 127.31 |
| Rpap1 | 7.69E-15 | 238.91 | 7.64E-10 | 120.42 | 5.60E-08 | 112.73 |
| Cul1 | 8.12E-11 | 236.19 | 0.02 | 105.28 | 2.14E-14 | 121.06 |
| Ckap2l | 1.42E-11 | 234.60 | 0.00 | 109.48 | 2.20E-16 | 142.69 |
| Mrpl22 | 9.05E-15 | 234.45 | 4.00E-06 | 112.30 | 0.16 | 103.22 |
| Bola3 | 1.60E-06 | 234.02 | 0.00 | 109.75 | 2.20E-16 | 138.64 |
| Rufy3 | 1.11E-09 | 233.93 | 0.02 | 93.57 | 7.37E-16 | 123.18 |
| Gm5089 | 1.75E-12 | 233.92 | 0.14 | 104.54 | 5.44E-15 | 127.90 |
| Tmem47 | 1.54E-12 | 233.42 | 2.49E-07 | 129.11 | 3.56E-16 | 126.88 |
| Ttll12 | 7.06E-07 | 232.20 | 0.01 | 108.55 | 2.20E-16 | 145.75 |
| Caml | 1.51E-11 | 231.77 | 0.84 | 99.07 | 1.13E-14 | 123.27 |
| Vmn1r43 | 1.05E-10 | 230.54 | 7.30E-08 | 119.28 | 2.20E-16 | 130.60 |
| Aldh16a1 | 8.22E-15 | 230.30 | 3.16E-13 | 124.11 | 0.00 | 107.76 |
| Gjb5 | 1.37E-06 | 228.50 | 0.02 | 109.01 | 1.54E-14 | 134.06 |
| Mum1 | 4.54E-14 | 228.00 | 0.00 | 109.40 | 3.20E-16 | 126.19 |
| Fgf2 | 3.56E-10 | 227.57 | 0.09 | 102.42 | 4.81E-13 | 123.45 |
| Dap3 | 1.14E-11 | 227.05 | 7.17E-06 | 114.62 | 0.00 | 107.35 |
| Cdkal1 | 7.41E-12 | 226.30 | 1.98E-06 | 88.53 | 3.30E-08 | 114.47 |
| Akap10 | 1.10E-11 | 226.09 | 3.08E-07 | 114.13 | 2.20E-16 | 133.61 |
| Dgka | 8.92E-14 | 225.20 | 0.04 | 106.02 | 2.14E-13 | 121.90 |
| Masp2 | 3.53E-09 | 225.18 | 3.32E-05 | 111.50 | 6.02E-07 | 111.19 |
| Mmgt1 | 1.18E-10 | 224.90 | 1.62E-12 | 124.16 | 1.01E-05 | 110.24 |
| Irf7 | 4.23E-06 | 224.54 | 0.41 | 104.58 | 0.00 | 106.65 |
| Lins | 8.51E-10 | 223.44 | 0.00 | 109.12 | 2.20E-16 | 132.09 |
| Pdcd6 | 1.53E-13 | 223.34 | 0.10 | 105.46 | 1.35E-12 | 123.57 |
| 4933425O20Rik | 1.74E-14 | 221.19 | 2.09E-15 | 129.67 | 2.25E-08 | 113.93 |
| Cd248 | 3.97E-12 | 220.04 | 3.18E-10 | 120.37 | 2.03E-15 | 127.05 |
| Scube3 | 6.66E-06 | 219.78 | 0.00 | 108.60 | 3.66E-14 | 132.37 |
| Clec4e | 1.33E-08 | 219.18 | 0.21 | 103.71 | 2.20E-16 | 133.57 |
| Tnfrsf26 | 8.83E-14 | 217.91 | 0.02 | 106.21 | 2.20E-16 | 142.27 |
| Wdr8 | 9.40E-15 | 216.91 | 5.14E-10 | 117.41 | 1.01E-15 | 129.06 |
| Kiss1r | 1.33E-13 | 216.76 | 0.39 | 97.76 | 5.82E-13 | 119.02 |
| Nudcd2 | 6.28E-10 | 216.74 | 0.05 | 106.28 | 0.05 | 104.80 |
| Gpr135 | 1.06E-09 | 216.54 | 9.46E-05 | 109.50 | 5.59E-15 | 123.38 |
| Tdrd7 | 2.64E-07 | 216.27 | 0.78 | 99.20 | 7.64E-15 | 129.02 |
| Chic1 | 2.85E-13 | 215.72 | 0.00 | 115.87 | 2.58E-15 | 126.13 |
| Tlr8 | 5.83E-11 | 215.20 | 0.00 | 108.48 | 1.80E-13 | 121.98 |
| D19Wsu162e | 5.41E-14 | 214.93 | 2.87E-10 | 118.31 | 1.52E-11 | 118.83 |
| Bdh2 | 8.92E-10 | 212.16 | 0.00 | 106.84 | 7.18E-11 | 116.74 |
| Hs3st1 | 1.73E-13 | 212.02 | 1.81E-09 | 119.15 | 2.24E-13 | 121.89 |
| Ptgr1 | 6.29E-12 | 211.36 | 3.77E-05 | 111.21 | 5.02E-13 | 121.05 |
| Arx | 1.57E-11 | 210.95 | 9.47E-05 | 110.99 | 6.37E-13 | 122.32 |
| Cc2d2a | 7.03E-14 | 209.96 | 1.38E-05 | 112.50 | 2.20E-16 | 130.71 |
| Pcgf3 | 1.34E-09 | 209.77 | 0.62 | 98.79 | 1.76E-14 | 125.62 |
| Chst14 | 9.16E-09 | 209.44 | 0.67 | 100.04 | 6.50E-14 | 123.98 |
| Zfp101 | 3.49E-13 | 207.42 | 0.02 | 106.45 | 2.20E-16 | 131.29 |
| Itgb4 | 4.30E-11 | 207.41 | 0.04 | 105.38 | 3.82E-14 | 121.99 |
| Tmem126a | 5.76E-14 | 206.82 | 1.30E-06 | 111.71 | 2.20E-16 | 128.10 |
| Maged1 | 1.21E-12 | 206.30 | 0.02 | 104.59 | 2.64E-14 | 120.52 |
| Il1rl2 | 1.50E-09 | 206.09 | 0.57 | 101.63 | 1.42E-13 | 124.01 |
| Ccdc50 | 3.20E-12 | 205.41 | 0.77 | 101.20 | 3.14E-15 | 124.27 |
| Srp9 | 7.02E-13 | 203.99 | 0.12 | 103.84 | 2.20E-16 | 125.95 |
| 2510006D16Rik | 2.65E-10 | 203.06 | 0.23 | 96.92 | 1.66E-14 | 125.41 |
| Plp2 | 9.83E-12 | 202.56 | 1.68E-06 | 114.97 | 2.39E-11 | 119.54 |
| Mael | 1.86E-09 | 202.32 | 6.35E-05 | 113.28 | 1.12E-11 | 118.24 |
| Chuk | 3.35E-10 | 201.89 | 0.00 | 107.80 | 0.66 | 100.27 |
| Gapvd1 | 8.21E-10 | 201.79 | 5.18E-06 | 112.49 | 4.79E-11 | 122.60 |
| Mon1a | 1.53E-08 | 201.21 | 2.56E-09 | 116.74 | 0.96 | 100.43 |
| Fbxo9 | 1.33E-10 | 200.50 | 0.00 | 109.60 | 5.06E-06 | 111.77 |
| Fam136a | 2.09E-10 | 198.72 | 0.00 | 110.83 | 4.29E-07 | 111.56 |
| Tgfb3 | 1.17E-10 | 198.32 | 0.19 | 97.46 | 3.26E-10 | 117.66 |
| Mmadhc | 2.77E-12 | 198.30 | 0.02 | 104.83 | 2.20E-16 | 126.17 |

| | | | | | | |
|---|---|---|---|---|---|---|
| Nfkbia | 9.99E-14 | 197.96 | 0.58 | 100.75 | 7.18E-15 | 126.20 |
| Nudt3 | 1.34E-11 | 197.40 | 0.02 | 105.79 | 2.20E-16 | 127.21 |
| Eps15 | 1.40E-10 | 196.67 | 1.09E-06 | 112.99 | 4.51E-11 | 116.98 |
| Ttll9 | 2.28E-05 | 194.84 | 5.88E-10 | 119.65 | 0.70 | 100.69 |
| Stk11 | 2.58E-09 | 193.34 | 0.02 | 106.17 | 8.00E-13 | 121.24 |
| Bbs10 | 1.89E-08 | 193.07 | 6.67E-07 | 114.47 | 0.00 | 92.19 |
| Cnnm4 | 7.07E-12 | 192.83 | 0.15 | 103.62 | 1.19E-14 | 124.09 |
| Pskh1 | 1.64E-07 | 192.58 | 0.17 | 97.27 | 8.27E-07 | 111.37 |
| Dclre1b | 6.98E-10 | 191.54 | 0.00 | 91.71 | 4.17E-16 | 123.05 |
| 4930590J08Rik | 1.89E-08 | 191.52 | 2.32E-05 | 90.00 | 5.51E-05 | 110.82 |
| Blnk | 0.00 | 191.29 | 0.00 | 94.01 | 5.24E-06 | 89.09 |
| Mzt1 | 2.41E-08 | 190.96 | 0.07 | 95.79 | 6.05E-14 | 122.68 |
| Slc38a3 | 3.24E-06 | 190.90 | 0.00 | 108.84 | 0.01 | 93.27 |
| Clpb | 1.15E-08 | 190.06 | 3.25E-06 | 114.10 | 1.38E-08 | 114.71 |
| Myocd | 9.99E-12 | 189.54 | 0.01 | 108.59 | 4.55E-09 | 116.03 |
| Pram1 | 7.29E-09 | 189.41 | 0.00 | 110.61 | 8.20E-13 | 120.51 |
| Snx16 | 2.51E-05 | 188.92 | 0.90 | 100.14 | 2.96E-16 | 130.89 |
| Cdkn1a | 6.44E-09 | 188.46 | 4.22E-12 | 122.56 | 2.20E-16 | 62.96 |
| Cfh | 1.13E-08 | 188.34 | 1.19E-06 | 89.40 | 1.91E-10 | 115.91 |
| Cpeb4 | 1.10E-10 | 188.27 | 0.08 | 105.39 | 1.53E-11 | 116.17 |
| Pak1ip1 | 1.75E-12 | 187.84 | 9.84E-10 | 116.90 | 0.00 | 107.83 |
| Unc45a | 1.68E-10 | 186.01 | 0.00 | 108.44 | 1.17E-12 | 79.20 |
| Arhgap23 | 5.91E-11 | 185.84 | 0.04 | 103.90 | 1.31E-12 | 119.98 |
| Rhbdl3 | 2.29E-09 | 182.95 | 0.50 | 98.85 | 9.06E-13 | 127.75 |
| Tgfbr3 | 1.30E-09 | 182.53 | 0.00 | 106.43 | 1.26E-12 | 120.42 |
| Mas1 | 4.80E-09 | 182.39 | 0.45 | 98.47 | 2.20E-16 | 124.31 |
| Camkk2 | 2.64E-07 | 181.83 | 0.08 | 105.43 | 0.17 | 102.34 |
| Hmgcll1 | 2.00E-07 | 181.52 | 0.40 | 99.65 | 1.69E-13 | 125.16 |
| Fabp3 | 1.01E-09 | 179.79 | 1.34E-06 | 112.38 | 4.04E-05 | 109.44 |
| Acp2 | 1.27E-08 | 179.65 | 0.42 | 102.51 | 9.37E-10 | 116.83 |
| Tmem128 | 4.92E-09 | 179.07 | 0.01 | 93.77 | 1.05E-07 | 112.22 |
| Nt5c | 2.27E-05 | 178.19 | 0.06 | 95.51 | 2.09E-10 | 118.00 |
| Rnf130 | 2.55E-08 | 177.38 | 1.69E-07 | 88.06 | 3.68E-11 | 116.82 |
| Smg6 | 6.28E-09 | 177.29 | 2.63E-15 | 76.95 | 0.04 | 124.29 |
| Aqp9 | 7.25E-08 | 177.00 | 0.73 | 100.57 | 7.78E-10 | 115.65 |
| Cr1l | 1.33E-07 | 174.85 | 0.00 | 91.70 | 0.08 | 103.29 |
| Rapgef4 | 1.07E-06 | 173.99 | 0.79 | 98.65 | 2.34E-10 | 117.74 |
| Btbd17 | 9.27E-07 | 173.71 | 0.14 | 103.95 | 1.03E-07 | 111.74 |
| Fads1 | 3.59E-09 | 173.25 | 0.53 | 98.49 | 0.00 | 107.46 |
| Idh3a | 6.47E-08 | 173.16 | 0.98 | 100.91 | 2.41E-10 | 117.84 |
| Muted | 3.02E-07 | 172.42 | 0.19 | 96.72 | 0.08 | 94.51 |
| Gsto1 | 8.41E-08 | 172.27 | 0.02 | 107.12 | 7.54E-07 | 111.63 |
| Siglece | 8.20E-05 | 172.16 | 0.13 | 95.97 | 5.70E-12 | 119.72 |
| Atg14 | 3.45E-06 | 171.65 | 0.00 | 108.54 | 0.01 | 106.93 |
| Mknk2 | 5.19E-05 | 170.96 | 1.14E-08 | 85.80 | 2.63E-15 | 124.61 |
| 2610301B20Rik | 8.37E-07 | 169.43 | 0.77 | 100.59 | 7.77E-08 | 111.88 |
| Man2b1 | 6.93E-09 | 168.06 | 9.77E-05 | 91.26 | 2.82E-08 | 113.47 |
| Ptcd1 | 1.85E-08 | 167.17 | 0.51 | 102.30 | 0.00 | 107.68 |
| Fbxl3 | 8.38E-09 | 166.61 | 0.01 | 107.39 | 0.25 | 103.23 |
| Dmrt2 | 5.63E-08 | 166.15 | 0.41 | 101.54 | 6.42E-05 | 90.60 |
| Tmem35 | 2.22E-08 | 164.93 | 0.08 | 95.77 | 0.00 | 107.30 |
| Tex19.1 | 4.58E-08 | 164.67 | 6.23E-08 | 112.55 | 1.21E-08 | 86.72 |
| Arl4c | 2.42E-05 | 164.31 | 0.09 | 104.42 | 1.91E-06 | 110.43 |
| Map3k2 | 2.36E-05 | 164.01 | 0.80 | 101.31 | 0.27 | 102.75 |
| Iffo2 | 2.72E-05 | 163.50 | 0.04 | 94.87 | 0.08 | 104.63 |
| Drg2 | 1.83E-07 | 161.47 | 0.12 | 103.72 | 0.84 | 99.54 |
| Ak4 | 2.46E-05 | 160.36 | 0.06 | 105.65 | 3.24E-11 | 81.63 |
| Reep1 | 1.26E-08 | 159.79 | 0.06 | 95.21 | 0.09 | 95.82 |
| Mapkapk2 | 5.41E-06 | 159.12 | 0.65 | 101.06 | 0.00 | 92.54 |
| Fbxo18 | 1.31E-05 | 157.98 | 0.24 | 103.29 | 0.79 | 100.80 |
| Atg4c | 1.75E-07 | 157.51 | 0.15 | 96.09 | 0.00 | 90.41 |
| Olfr986 | 4.57E-08 | 157.03 | 0.00 | 91.86 | 0.01 | 105.07 |
| Calcrl | 0.00 | 154.85 | 0.02 | 106.39 | 0.08 | 95.27 |
| Tgm2 | 7.88E-05 | 154.75 | 1.15E-07 | 85.95 | 9.49E-14 | 124.90 |
| 2610528E23Rik | 1.38E-08 | 153.73 | 0.00 | 108.57 | 8.09E-06 | 89.84 |
| 2700049A03Rik | 9.00E-07 | 152.25 | 2.45E-12 | 82.28 | 3.79E-10 | 114.43 |

| | | | | | | |
|---|---|---|---|---|---|---|
| Zfyve1 | 1.04E-06 | 151.14 | 2.28E-05 | 111.79 | 0.02 | 94.37 |
| Efhc1 | 9.54E-06 | 147.46 | 6.28E-07 | 88.09 | 0.88 | 99.69 |
| Tas2r126 | 0.00 | 145.69 | 1.30E-07 | 86.36 | 4.52E-09 | 85.10 |
| Mthfd2 | 0.00 | 145.20 | 0.15 | 103.75 | 0.18 | 94.83 |
| Myo1e | 1.91E-05 | 141.26 | 0.77 | 100.14 | 0.17 | 96.14 |
