## Supplementary material for "Auto-inhibition of myoblast fusion by cyclic receptor signalling": Table 2

### Table 2. Genes necessary for C2C12 fusion

| Entrez Gene ID | pvalue | %nuclei/myotube | pvalue | %nuclei | pvalue | %myogenin |
|---|---|---|---|---|---|---|
| Mrps10 | 2.20E-16 | 34.13 | 1.20E-15 | 77.32 | 3.89E-12 | 76.46 |
| Tfrc | 2.20E-16 | 34.95 | 2.20E-16 | 63.29 | 2.20E-16 | 69.01 |
| Fau | 2.20E-16 | 38.18 | 3.15E-16 | 77.16 | 7.78E-14 | 77.76 |
| Psmd6 | 2.20E-16 | 39.28 | 5.92E-16 | 74.56 | 2.20E-16 | 66.09 |
| Snrpd1 | 2.20E-16 | 40.39 | 2.20E-16 | 74.32 | 1.58E-11 | 79.81 |
| Eif2s1 | 2.20E-16 | 44.04 | 3.26E-11 | 82.37 | 0.01 | 92.34 |
| Prpf8 | 2.20E-16 | 44.35 | 2.24E-06 | 87.23 | 1.07E-09 | 85.17 |
| Sdad1 | 2.20E-16 | 44.99 | 2.20E-16 | 72.94 | 0.02 | 94.73 |
| Dcaf13 | 2.20E-16 | 45.03 | 2.48E-14 | 76.98 | 9.57E-10 | 84.08 |
| Mprip | 2.13E-15 | 45.33 | 5.95E-09 | 84.22 | 4.79E-13 | 77.27 |
| 9830001H06Rik | 2.20E-16 | 45.33 | 0.01 | 104.89 | 1.66E-06 | 90.10 |
| Klhdc3 | 2.20E-16 | 46.10 | 0.67 | 100.54 | 1.42E-15 | 76.26 |
| Hmx1 | 2.20E-16 | 46.15 | 8.88E-08 | 87.17 | 3.13E-14 | 80.67 |
| Sec24a | 2.20E-16 | 46.51 | 4.22E-08 | 86.35 | 3.47E-13 | 81.26 |
| Atp7b | 5.72E-16 | 46.91 | 0.01 | 93.72 | 3.38E-08 | 86.39 |
| Hook3 | 2.07E-15 | 47.42 | 1.44E-13 | 81.99 | 2.20E-16 | 72.09 |
| Mapk14 | 1.10E-14 | 47.61 | 2.63E-09 | 86.96 | 2.20E-16 | 71.97 |
| Edf1 | 1.06E-15 | 48.02 | 0.01 | 107.11 | 2.20E-16 | 57.52 |
| Olfr1321 | 7.73E-15 | 48.18 | 7.56E-14 | 79.04 | 2.01E-06 | 87.60 |
| Ubap2l | 2.20E-16 | 48.35 | 1.04E-11 | 83.36 | 6.74E-06 | 87.17 |
| Zfp513 | 2.04E-14 | 48.50 | 4.44E-12 | 80.91 | 3.15E-09 | 83.70 |
| Tmem40 | 1.85E-15 | 48.56 | 0.86 | 99.76 | 3.50E-16 | 70.35 |
| AA467197 | 8.38E-15 | 48.73 | 0.00 | 107.42 | 1.24E-11 | 82.15 |
| Ubxn10 | 1.48E-14 | 48.78 | 0.00 | 89.80 | 0.00 | 90.70 |
| Gnl3l | 1.50E-15 | 48.95 | 7.69E-15 | 73.72 | 0.00 | 85.27 |
| Micalcl | 2.25E-15 | 49.06 | 3.09E-10 | 82.43 | 0.00 | 108.67 |
| Plcg2 | 7.19E-16 | 49.09 | 8.74E-11 | 83.13 | 5.45E-13 | 82.29 |
| Rgmb | 2.20E-16 | 49.23 | 0.45 | 101.30 | 0.46 | 99.16 |
| Katnb1 | 1.56E-15 | 49.23 | 4.16E-08 | 86.79 | 5.76E-06 | 89.17 |
| Prrc2c | 2.18E-13 | 49.54 | 2.93E-15 | 75.56 | 0.22 | 96.23 |
| Gsk3b | 1.94E-15 | 49.62 | 3.13E-06 | 88.71 | 6.85E-14 | 77.28 |
| Asap1 | 3.32E-14 | 50.45 | 1.12E-08 | 85.22 | 4.55E-15 | 77.80 |
| Dhx30 | 1.48E-14 | 50.61 | 2.30E-08 | 84.81 | 0.00 | 107.52 |
| Pknox1 | 4.13E-15 | 50.90 | 9.86E-14 | 78.18 | 0.00 | 90.17 |
| Blm | 1.38E-13 | 50.91 | 1.37E-06 | 87.50 | 3.83E-12 | 79.99 |
| Bnip2 | 4.29E-14 | 51.39 | 5.46E-08 | 85.91 | 3.93E-05 | 89.26 |
| Exoc6b | 4.75E-13 | 52.05 | 0.02 | 106.49 | 2.07E-06 | 89.12 |
| Plekhf1 | 1.83E-12 | 52.39 | 9.24E-09 | 84.48 | 4.64E-11 | 79.86 |
| Catsper2 | 4.96E-14 | 52.48 | 0.01 | 93.97 | 0.64 | 98.39 |
| Dnalc4 | 1.24E-14 | 52.79 | 1.88E-06 | 89.89 | 1.28E-07 | 88.26 |
| Lingo2 | 3.43E-13 | 53.13 | 4.18E-07 | 87.23 | 0.00 | 91.84 |
| C1ql3 | 1.33E-12 | 53.19 | 6.11E-10 | 83.84 | 0.00 | 88.93 |
| Eci2 | 4.98E-13 | 53.24 | 0.40 | 101.42 | 0.50 | 99.18 |
| Atp6ap1 | 5.08E-13 | 53.75 | 9.39E-10 | 84.93 | 2.39E-07 | 113.69 |
| Arhgap30 | 1.49E-13 | 53.79 | 0.77 | 98.65 | 3.90E-07 | 113.62 |
| Crls1 | 5.77E-14 | 54.44 | 2.82E-05 | 110.84 | 1.10E-07 | 114.62 |
| Brca2 | 2.60E-12 | 54.49 | 3.92E-11 | 83.04 | 0.00 | 92.61 |
| Melk | 2.23E-14 | 54.69 | 0.04 | 95.16 | 8.23E-09 | 86.99 |
| Eci1 | 2.90E-14 | 55.44 | 0.30 | 97.41 | 1.06E-08 | 112.49 |
| Olfr1040 | 1.82E-12 | 55.71 | 2.18E-09 | 85.89 | 3.00E-05 | 88.05 |
| Amz1 | 4.71E-12 | 55.96 | 0.07 | 95.25 | 2.23E-07 | 88.52 |
| Tmem87a | 1.03E-12 | 56.51 | 2.23E-14 | 77.29 | 0.00 | 93.07 |
| Prkca | 2.61E-13 | 56.62 | 1.22E-09 | 84.68 | 2.80E-09 | 86.59 |
| Atp6v0e | 1.55E-11 | 56.92 | 4.11E-08 | 87.00 | 1.97E-06 | 89.53 |
| Sema5b | 5.62E-12 | 57.00 | 3.22E-10 | 84.15 | 2.86E-10 | 84.13 |
| Srsf1 | 7.37E-12 | 57.25 | 1.72E-10 | 83.92 | 0.02 | 106.10 |
| Arhgap17 | 2.63E-13 | 57.38 | 0.00 | 92.92 | 0.01 | 105.00 |
| Jarid2 | 1.57E-13 | 57.40 | 0.37 | 97.24 | 8.98E-10 | 83.56 |
| Tmem186 | 4.23E-11 | 57.54 | 0.02 | 106.17 | 0.00 | 106.47 |
| Trim45 | 3.97E-12 | 57.64 | 2.97E-09 | 85.88 | 0.88 | 99.66 |
| Idua | 3.27E-10 | 57.85 | 2.56E-11 | 80.31 | 3.41E-14 | 124.50 |
| 1700001O22Rik | 4.91E-09 | 58.19 | 0.97 | 100.94 | 0.00 | 88.76 |
| Gria4 | 5.67E-10 | 58.21 | 0.24 | 97.70 | 1.06E-14 | 75.38 |
| Cct8l1 | 3.11E-11 | 58.38 | 1.75E-08 | 84.59 | 5.27E-09 | 86.07 |

| | | | | | | |
|---|---|---|---|---|---|---|
| Triobp | 2.37E-12 | 59.10 | 6.77E-07 | 89.98 | 1.66E-13 | 81.21 |
| 1110008J03Rik | 1.14E-11 | 59.46 | 0.02 | 93.29 | 2.78E-13 | 78.42 |
| Rinl | 1.08E-11 | 59.63 | 0.00 | 91.35 | 8.60E-07 | 89.17 |
| Specc1l | 5.92E-11 | 59.87 | 0.11 | 104.26 | 0.93 | 99.75 |
| Pkd1l2 | 4.30E-10 | 60.41 | 1.87E-08 | 84.77 | 0.41 | 101.97 |
| Taf15 | 3.04E-11 | 61.06 | 0.00 | 91.21 | 0.17 | 97.18 |
| Chmp4c | 2.89E-09 | 61.11 | 0.07 | 105.50 | 1.00E-06 | 88.23 |
| Gpr30 | 9.29E-11 | 61.23 | 0.11 | 96.80 | 6.05E-12 | 81.99 |
| Zfp697 | 3.92E-10 | 61.33 | 7.61E-06 | 88.81 | 5.54E-06 | 86.81 |
| Smpd4 | 1.02E-10 | 61.70 | 0.01 | 106.34 | 0.00 | 94.52 |
| Creb3l4 | 2.83E-10 | 61.71 | 0.49 | 101.58 | 4.35E-12 | 80.25 |
| Cep97 | 1.21E-10 | 62.00 | 1.14E-06 | 88.62 | 8.24E-06 | 89.03 |
| Slc39a1 | 5.36E-10 | 62.04 | 0.09 | 95.26 | 0.17 | 97.23 |
| Sall4 | 2.28E-09 | 62.05 | 0.29 | 95.87 | 6.24E-13 | 80.52 |
| Ruvbl2 | 1.67E-09 | 62.20 | 0.46 | 98.00 | 6.21E-07 | 88.61 |
| Haus8 | 2.53E-09 | 62.21 | 0.53 | 100.78 | 0.00 | 92.19 |
| Lrmp | 3.37E-09 | 62.21 | 0.05 | 95.52 | 9.58E-09 | 85.86 |
| Dnajc24 | 2.39E-10 | 62.22 | 2.22E-09 | 85.50 | 0.01 | 94.89 |
| Psrc1 | 3.95E-10 | 62.40 | 1.15E-06 | 89.58 | 0.61 | 101.64 |
| Rgs5 | 2.05E-10 | 62.65 | 0.00 | 110.90 | 0.00 | 92.95 |
| Nlgn2 | 4.48E-10 | 62.78 | 0.00 | 92.14 | 0.00 | 92.54 |
| Hoxc5 | 9.63E-10 | 63.07 | 0.47 | 97.52 | 6.44E-12 | 82.57 |
| Ezr | 5.81E-09 | 63.19 | 0.02 | 94.03 | 2.45E-05 | 88.73 |
| Fktn | 3.35E-09 | 63.19 | 4.51E-06 | 87.41 | 0.66 | 100.94 |
| Kif3c | 1.58E-09 | 63.26 | 0.06 | 95.91 | 9.46E-11 | 84.46 |
| Ncoa2 | 1.95E-09 | 63.33 | 0.00 | 92.57 | 2.22E-08 | 86.14 |
| Wwtr1 | 1.39E-10 | 63.47 | 0.01 | 92.94 | 6.57E-14 | 120.73 |
| Hmbs | 2.53E-09 | 63.71 | 8.05E-14 | 127.11 | 2.58E-05 | 90.81 |
| Zdhhc25 | 3.48E-09 | 63.74 | 0.00 | 106.43 | 1.27E-06 | 111.36 |
| Fam125a | 6.60E-10 | 63.77 | 9.67E-08 | 111.96 | 2.76E-12 | 81.19 |
| Bag3 | 7.84E-10 | 63.85 | 0.00 | 107.11 | 6.55E-07 | 88.63 |
| Itgae | 2.18E-09 | 64.04 | 0.00 | 108.79 | 7.97E-07 | 88.10 |
| Mepce | 2.16E-08 | 64.05 | 0.10 | 103.55 | 2.49E-06 | 110.18 |
| Irak1 | 8.26E-10 | 64.19 | 2.22E-10 | 118.00 | 1.55E-11 | 84.00 |
| Tspan7 | 1.19E-09 | 64.24 | 0.54 | 98.36 | 0.13 | 97.09 |
| Rcsd1 | 3.52E-09 | 64.31 | 0.31 | 97.48 | 4.71E-07 | 112.66 |
| Dennd4b | 1.82E-08 | 64.43 | 0.00 | 92.78 | 0.71 | 101.01 |
| Idh1 | 7.06E-09 | 64.76 | 3.99E-05 | 89.52 | 0.01 | 104.90 |
| Psmc5 | 2.51E-08 | 64.78 | 0.56 | 101.33 | 0.00 | 91.91 |
| Zfp341 | 3.49E-09 | 64.79 | 0.00 | 91.49 | 0.01 | 94.48 |
| Dnajb5 | 1.81E-08 | 65.18 | 0.35 | 97.96 | 0.01 | 94.72 |
| Fgf3 | 8.80E-09 | 65.32 | 5.90E-09 | 120.01 | 0.01 | 105.52 |
| Rab39 | 5.48E-08 | 65.33 | 0.39 | 101.87 | 0.01 | 93.02 |
| Tcf15 | 3.37E-09 | 65.46 | 0.00 | 108.27 | 1.60E-06 | 87.42 |
| Ppp2r1a | 2.70E-08 | 65.56 | 0.08 | 95.76 | 3.12E-05 | 90.72 |
| Cnot6l | 9.02E-08 | 65.57 | 0.22 | 102.16 | 0.02 | 95.61 |
| Gm2a | 2.54E-08 | 65.61 | 0.58 | 98.74 | 0.00 | 91.82 |
| Srrm2 | 2.30E-08 | 65.70 | 0.87 | 99.24 | 0.02 | 94.13 |
| Tinf2 | 2.81E-08 | 65.89 | 0.02 | 94.45 | 0.23 | 102.17 |
| Cldn13 | 1.31E-08 | 65.95 | 3.35E-10 | 121.81 | 4.20E-06 | 109.84 |
| 1810037I17Rik | 3.54E-08 | 66.07 | 0.13 | 96.56 | 8.33E-09 | 85.14 |
| Syt7 | 3.68E-09 | 66.16 | 0.00 | 105.22 | 0.00 | 92.93 |
| Vta1 | 5.68E-08 | 66.18 | 0.09 | 96.15 | 0.00 | 92.64 |
| Reep2 | 1.71E-07 | 66.27 | 0.00 | 91.91 | 0.01 | 94.36 |
| Galntl6 | 1.87E-07 | 66.38 | 3.53E-12 | 80.75 | 0.01 | 105.62 |
| Rhod | 1.35E-07 | 66.42 | 0.00 | 93.24 | 0.00 | 108.82 |
| Cpsf1 | 6.47E-08 | 66.54 | 1.26E-05 | 111.04 | 0.04 | 96.26 |
| Zbtb46 | 1.65E-08 | 66.81 | 4.57E-09 | 86.19 | 0.56 | 100.76 |
| Apba1 | 9.07E-08 | 66.86 | 0.02 | 95.05 | 0.37 | 103.25 |
| Syt5 | 2.79E-08 | 66.95 | 0.04 | 95.06 | 0.03 | 94.98 |
| Grpel1 | 1.45E-07 | 67.09 | 0.35 | 101.64 | 3.76E-06 | 89.89 |
| Metap2 | 8.05E-07 | 67.12 | 0.03 | 92.58 | 0.09 | 95.98 |
| Cenpb | 6.87E-08 | 67.12 | 0.03 | 95.33 | 0.22 | 97.58 |
| Bcor | 4.40E-08 | 67.15 | 5.77E-07 | 88.41 | 0.31 | 98.01 |
| Usp29 | 1.17E-07 | 67.68 | 0.09 | 96.18 | 0.00 | 108.15 |
| Treml1 | 4.17E-07 | 67.82 | 0.26 | 102.26 | 0.56 | 98.89 |

| | | | | | | |
|---|---|---|---|---|---|---|
| Fxyd5 | 1.10E-07 | 68.03 | 4.03E-05 | 89.70 | 0.15 | 96.18 |
| Kcnh3 | 7.05E-07 | 68.05 | 0.00 | 91.25 | 0.12 | 97.07 |
| Aoc3 | 1.76E-07 | 68.06 | 0.00 | 109.96 | 8.57E-07 | 113.09 |
| Fhl1 | 2.03E-06 | 68.09 | 0.00 | 110.02 | 2.01E-08 | 84.38 |
| Ncapd3 | 3.25E-07 | 68.45 | 0.79 | 98.62 | 2.25E-07 | 113.36 |
| Plat | 2.39E-06 | 68.49 | 0.98 | 100.83 | 6.62E-05 | 112.13 |
| Slc26a11 | 1.68E-06 | 68.61 | 0.07 | 107.08 | 0.01 | 106.09 |
| Stxbp1 | 4.36E-07 | 68.62 | 0.00 | 108.28 | 0.00 | 92.18 |
| Hook2 | 8.68E-07 | 68.77 | 2.20E-16 | 143.06 | 0.65 | 101.04 |
| 6030419C18Rik | 3.82E-07 | 69.00 | 0.74 | 98.46 | 0.01 | 93.50 |
| Cdh15 | 6.29E-07 | 69.01 | 0.00 | 107.96 | 0.42 | 101.93 |
| Pi4k2b | 2.32E-07 | 69.22 | 0.45 | 98.22 | 1.19E-07 | 111.81 |
| Skor1 | 6.04E-07 | 69.23 | 0.61 | 101.54 | 0.01 | 93.59 |
| Runx1t1 | 3.13E-06 | 69.47 | 0.97 | 99.53 | 4.39E-05 | 91.55 |
| Rin1 | 8.89E-07 | 70.21 | 0.24 | 96.03 | 3.23E-15 | 125.35 |
| Ccdc104 | 1.29E-05 | 70.33 | 8.52E-05 | 88.70 | 3.35E-06 | 114.59 |
| Fgf15 | 3.60E-06 | 70.93 | 0.69 | 100.46 | 2.12E-05 | 89.07 |
| Col8a1 | 7.17E-06 | 70.96 | 0.14 | 95.64 | 0.78 | 100.54 |
| Plk2 | 4.31E-06 | 71.02 | 0.00 | 92.16 | 0.11 | 103.68 |
| Ppt2 | 2.98E-06 | 71.04 | 0.00 | 93.22 | 0.00 | 108.02 |
| Mcam | 2.34E-06 | 71.08 | 4.20E-06 | 110.29 | 0.00 | 93.40 |
| Naca | 1.42E-05 | 71.12 | 0.24 | 103.05 | 0.00 | 89.68 |
| Chchd7 | 6.73E-06 | 71.20 | 3.41E-06 | 89.78 | 0.48 | 97.96 |
| Egfr | 3.54E-06 | 71.21 | 0.46 | 98.36 | 0.01 | 105.14 |
| Cxcr7 | 2.46E-06 | 71.58 | 0.44 | 101.16 | 0.00 | 105.50 |
| A2m | 9.62E-06 | 71.69 | 0.04 | 95.46 | 0.00 | 106.53 |
| Gja4 | 3.88E-05 | 71.81 | 0.75 | 103.28 | 2.01E-05 | 89.98 |
| Fam111a | 2.25E-05 | 71.87 | 0.48 | 97.97 | 0.83 | 100.31 |
| Syp | 3.83E-06 | 71.89 | 0.25 | 103.19 | 3.43E-13 | 119.78 |
| Stag2 | 5.85E-06 | 72.02 | 4.21E-09 | 85.46 | 0.14 | 103.32 |
| Psmc2 | 3.09E-05 | 72.05 | 0.03 | 96.53 | 0.00 | 93.66 |
| 8430408G22Rik | 1.60E-05 | 72.08 | 0.14 | 103.00 | 0.15 | 96.73 |
| 2310043J07Rik | 5.77E-06 | 72.14 | 0.00 | 92.20 | 0.60 | 101.36 |
| Ror2 | 3.20E-05 | 72.23 | 0.00 | 109.23 | 0.00 | 93.19 |
| Acads | 6.88E-05 | 72.25 | 0.00 | 91.41 | 0.00 | 109.84 |
| Armcx6 | 3.49E-05 | 72.30 | 0.66 | 99.13 | 0.03 | 105.83 |
| Nif3l1 | 0.00 | 72.52 | 1.35E-05 | 89.40 | 0.02 | 100.01 |
| Mpg | 9.30E-06 | 72.61 | 0.01 | 94.91 | 0.00 | 108.37 |
| Dido1 | 3.33E-05 | 72.69 | 0.23 | 103.46 | 7.03E-05 | 107.87 |
| Nckap1 | 7.67E-06 | 72.70 | 0.23 | 97.18 | 2.20E-16 | 130.30 |
| Klc4 | 1.60E-05 | 72.86 | 0.02 | 104.36 | 3.72E-05 | 87.60 |
| Gata4 | 1.24E-05 | 73.14 | 0.18 | 102.49 | 0.01 | 93.53 |
| Akr1c12 | 0.00 | 73.39 | 3.58E-05 | 90.11 | 0.01 | 105.24 |
| Marveld2 | 1.66E-05 | 73.40 | 1.33E-11 | 125.18 | 2.20E-16 | 138.46 |
| Aldoc | 5.43E-05 | 73.42 | 0.52 | 101.75 | 0.65 | 101.06 |
| Kank2 | 6.92E-05 | 73.70 | 0.05 | 105.10 | 5.52E-07 | 86.51 |
| Il4ra | 2.30E-07 | 73.74 | 0.00 | 106.98 | 0.04 | 96.08 |
| Col6a2 | 4.23E-05 | 73.75 | 0.00 | 110.30 | 2.51E-08 | 87.24 |
| Arhgef7 | 5.11E-05 | 73.80 | 0.01 | 93.60 | 0.08 | 103.46 |
| Ap2m1 | 6.34E-05 | 73.83 | 0.09 | 96.28 | 1.69E-07 | 115.15 |
| Fbl | 9.77E-05 | 73.84 | 0.05 | 95.59 | 0.00 | 107.64 |
| Cdc25b | 0.00 | 73.92 | 0.77 | 103.15 | 7.43E-07 | 88.43 |
| Fus | 4.89E-05 | 74.05 | 0.00 | 106.34 | 1.25E-06 | 109.31 |
| Ccl2 | 7.51E-06 | 74.51 | 0.25 | 96.21 | 0.95 | 99.76 |
| Exoc3l | 0.00 | 74.92 | 3.35E-05 | 109.53 | 0.76 | 98.97 |
| Rnf123 | 7.14E-05 | 75.05 | 0.64 | 100.97 | 4.26E-09 | 114.08 |
| Rcc2 | 0.00 | 75.34 | 1.66E-10 | 117.21 | 0.05 | 96.34 |
| Sphk1 | 0.00 | 75.56 | 9.54E-10 | 117.15 | 2.19E-09 | 86.44 |
| Lrtm2 | 0.00 | 75.62 | 0.41 | 101.97 | 0.47 | 102.04 |
| Scp2 | 0.00 | 76.24 | 9.34E-16 | 76.99 | 3.53E-14 | 121.93 |
| Rtn4ip1 | 0.00 | 76.30 | 0.84 | 100.23 | 0.80 | 100.70 |
| Dicer1 | 0.00 | 76.49 | 0.99 | 99.40 | 3.12E-08 | 111.89 |
| Arap3 | 0.00 | 76.54 | 0.00 | 93.38 | 0.00 | 106.59 |
| Tmem63a | 0.00 | 76.61 | 0.97 | 99.30 | 0.88 | 99.14 |
| Scaf1 | 0.00 | 77.13 | 0.01 | 108.58 | 0.00 | 105.95 |
| Zfp105 | 0.00 | 77.29 | 0.28 | 102.98 | 1.93E-06 | 112.93 |

| | | | | | | |
|---|---|---|---|---|---|---|
| Rtn2 | 0.00 | 77.31 | 0.26 | 95.97 | 0.00 | 108.61 |
| Pdcd4 | 0.00 | 77.49 | 0.01 | 106.45 | 4.05E-08 | 113.10 |
| Sox15 | 0.00 | 77.58 | 0.00 | 92.23 | 0.02 | 104.41 |
| Fermt3 | 0.00 | 79.35 | 7.73E-05 | 111.41 | 0.02 | 104.79 |
| 4930563D23Rik | 0.00 | 82.53 | 0.00 | 93.65 | 4.47E-09 | 118.02 |
