## Supplementary material for "Auto-inhibition of myoblast fusion by cyclic receptor signalling": Table 3

### Table 3. Role of TGFβ-associated genes on C2C12 fusion

| TGFβ ligand | Fusion (% of Ctrl) | Proliferation | Differentiation (MyoG ) |
|---|---|---|---|
| TGFB1 | ns | | |
| TGFB2 | 406 | 112 | 154 |
| TGFB3 | 198 | 97 | 118 |
| TGFβ receptor | | | |
| TGFBR1 | 284 | 97 | 149 |
| TGFBR3 | 183 | 106 | 120 |
| ACVR1B | 254 | 112 | 124 |
| Effectors | | | |
| SMAD2 | 166 | 91 | 131 |
| SMAD3 | 378 | 100 | 137 |
| SMAD4 | 524 | 98 | 157 |
| Synergizes with TGFβ(ref) | | | |
| DPT(45) | 148 | 113 | 113 |
| MMP14(46) | 154 | 90 | 126 |
| RUNX1(47) | 179 | 107 | 125 |
| SCUBE3(48) | 220 | 108 | 132 |
| TGFβ antagonists | | | |
| TGIF1 | 57 | 88 | 79 |
| BMP pathway | | | |
| BMPR1A | ns | | |
| BMP 1, 4, 7, 8 | ns | | |
| ACVR2A, 2B | ns | | |
| SMAD5 | ns | | |
| Activin / inhibin | | | |
| INHA | ns | | |
| INHBC | ns | | |
| INHBE | ns | | |
| Positive controls | | | |
| RAC1 | 73 | 85 | 142 |
| CDC42 | 52 | 81 | 140 |

ns: non significant
